# IRG1 controls host responses to restrict *Mycobacterium tuberculosis* infection

**DOI:** 10.1101/761551

**Authors:** Arnaud Machelart, Imène Belhaouane, Nathalie Deboosere, Isabelle Poncin, Jean-Paul Saint-André, Anne-Marie Pauwels, Ok-Ryul Song, Samuel Jouny, Carine Rouanet, Anaïs Poncet, Sabrina Marion, William Laine, Jérôme Kluza, Eric Muraille, Rudi Beyaert, Laleh Majlessi, Stéphane Canaan, Priscille Brodin, Eik Hoffmann

**Affiliations:** Univ. Lille, CNRS, INSERM, CHU Lille, Institut Pasteur de Lille, U1019 - UMR 9017 - CIIL - Center for Infection and Immunity of Lille, Lille, France; Aix-Marseille Univ., CNRS, UMR 7255 - LISM, IMM - FR3479, Marseille, France; University Hospital Center of Angers, Angers, France; Unit of Molecular Signal Transduction in Inflammation, VIB-UGent Center for Inflammation Research, Ghent, Belgium; Department of Biomedical Molecular Biology, Ghent University, Ghent, Belgium; Institut pour la Recherche sur le cancer de Lille (IRCL), Canther, INSERM UMR 9020 – UMR-S 1277, Lille, France; Unité de Recherche en Biologie des Microorganismes (URBM), NARILIS, University of Namur, Namur, Belgium; Université Libre de Bruxelles, Laboratoire de Parasitologie, and ULB Center for Research in Immunology (U-CRI), Gosselies, Belgium; Institut Pasteur – TheraVectys Joint Lab, Virology Department, Paris, France

**Keywords:** host-pathogen interactions, metabolic reprogramming, tuberculosis, lipid droplet, macrophage, dendritic cell

## Abstract

*Mycobacterium tuberculosis* (*Mtb*), the pathogen causing human tuberculosis, has evolved multiple strategies to successfully prevent clearance by immune cells and to establish dissemination and long-term survival in the host. The modulation of host immunity to maximize pathogen elimination while minimizing inflammation-mediated tissue damage may provide another tool to fight drug-resistant *Mtb* strains. Metabolic reprogramming of phagocytes can dramatically influence the intracellular colonization by *Mtb* and the key players involved in this process remain a matter of debate. Here, we demonstrate that aconitate decarboxylase 1 (Acod1; also known as immune-responsive gene 1, IRG1), which converts cis-aconitate into the metabolite itaconate, is a major player in controlling the acute phase of *Mtb* infection. Exposure of IRG1-deficient mice to a virulent *Mtb* strain (H37Rv) was lethal, while *M. bovis* BCG and the H37Ra attenuated *Mtb* strain induced neither lethality nor severe lung immunopathology. Lungs of IRG1-deficient mice infected by *Mtb* H37Rv displayed large areas of necrotizing granulomatous inflammation and neutrophil infiltration, accompanied by reduced levels of B and T lymphocytes and increased levels of alveolar and interstitial macrophage populations, compared to their wild type counterparts. Next, we show that IRG1, beyond its recruitment to *Mtb*-containing vacuoles, restricts *Mtb* replication and lipid droplets accumulation in phagocytes, hallmarks of a tight interplay between the bacillus and the host. Altogether, IRG1 confines the host response to create a favourable phagocytic environment for *Mtb* controlled intracellular replication.

## Introduction

Protective immunity of host cells during their infection by bacterial pathogens includes a broad variety of pathways and spatially regulated molecular players. Although the interplay between mechanisms of antimicrobial resistance and adapted tolerance of inflammatory responses is able to control infection, e.g. in the lung, several pathogens have evolved strategies to resist host defense and to persist for long time periods. *Mycobacterium tuberculosis* (*Mtb*), responsible for tuberculosis (TB) in humans, is transmitted by aerosol droplets followed by engulfment by alveolar macrophages and dendritic cells (DCs) in the lung. *Mtb* is able to evade different innate antimicrobial mechanisms of host cells and replicates intracellularly [1]. In addition, host adaptive immune responses are activated and slow down mycobacterial growth, but *Mtb* infection can also lead to chronic forms of TB. Therefore, TB remains a leading cause of death worldwide, responsible for an estimated 1.5 million deaths each year, together with a dramatic increase in the emergence of multidrug- and extensively drug-resistant *Mtb* strains [2]. While any organ in the body can be affected by *Mtb* infection, new infectious cycles are induced by transmission of pulmonary forms of the disease [2]. The live attenuated *Mycobacterium bovis* strain Bacillus Calmette-Guérin (BCG) is the only available vaccine against TB, but is not sufficiently successful in preventing active TB in adults. BCG generates prolonged antigen-responsive CD4 and CD8 T cell responses and remains the gold standard in animal vaccine studies [3].

Host-directed therapies (HDTs) against bacterial infections are in development that support elimination of mycobacteria by the host while reducing tissue damage induced by the infection [4]. Advances in the understanding of key players involved in immunometabolism shed light on the intimate link between metabolic states of immune cells and their specific functions during infection and inflammation [5, 6] and are increasingly applied for the development of HDTs against different infectious diseases, including TB [7]. The *Mtb*-infected host cell microenvironment is characterized by dysregulated immunoregulation pathways, for example, Th1/Th17 versus Th2 balance, regulatory T and suppressive myeloid cell populations and a shift from M1-like to M2-like polarized macrophages [1]. Recent studies have shown that changes in specific host metabolites can be mapped to cellular effector mechanisms and drive different inflammatory phenotypes of immune cells [8]. Itaconate, a host metabolite that is produced by different immune cell populations upon pro-inflammatory stimuli, such as LPS and type I and II interferons [9, 10], was also found in the lungs of *Mtb*-infected mice [11] and received increasing attention in recent years [12, 13]. Itaconate is generated from cis-aconitate in the tricarboxylic acid (TCA) cycle by the catalytic enzyme immune-responsive gene 1 (IRG1), also known as aconitate decarboxylase 1 (ACOD1) [14]. It has previously been shown that itaconate has antimicrobial activity by inhibiting isocitrate lyase [15], an enzyme of the glyoxylate shunt, which is present in most prokaryotes but absent in mammals. On the host side, itaconate was shown to affect major inflammatory pathways in immune cells by blocking succinate dehydrogenase [16], by controlling IL-1β expression and NLRP3 inflammasome activation [17, 18, 19], and by regulating HIF-1α activity, production of antimicrobial ROS and NO by Nrf2 activation [17, 20, 21]. In turn, itaconate was also recently shown to suppress M2 macrophage polarization [22]. Itaconate activities are able to influence host-pathogen interactions, as it was shown during infections by *Legionella pneumophila* [23, 24], *Pseudomonas aeruginosa* [25], Zika virus [26], *Francisella tularensis* [27], *Salmonella enterica* [28], *Staphylococcus aureus* [29] and *Brucella melitensis* or *B. abortus* [30, 31]. Concerning TB, Nair et al. showed that IRG1-deficient mice are highly susceptible to *Mtb* infection, while no aberrant phenotypes were found during influenza A virus or *Listeria monocytogenes* infection [32]. Their findings suggest that IRG1 expression in myeloid cells shape immunometabolic host responses by regulating neutrophil-dependent inflammation during *Mtb* infection of the lung. However, the underlying intracellular activities of *Mtb*-infected immune cells and their contribution to the observed phenotype remained unknown.

Similar to the previous report using aerosol infection [32], we show here that intranasal inoculation of IRG1-deficient mice by *Mtb* H37Rv induced severe lung immunopathology and mortality of infected mice. Exacerbated inflammation and high mycobacterial burden in the lungs of *Mtb*-infected, IRG1-deficient mice were accompanied by large areas of necrotizing granuloma formation, neutrophil infiltration and a pronounced reduction in the number of B and T lymphocytes. Interestingly, exposure of IRG1-deficient mice to the attenuated *Mtb* strain H37Ra or the vaccinal *M. bovis* BCG strain via the intranasal route induced neither lethality nor severe lung immunopathology demonstrating that the phenotype observed in *Mtb*-infected mice is linked to pathogenic virulence. Moreover, we show that IRG1 is induced upon *Mtb* infection and is directly recruited to *Mtb*-containing phagosomes. IRG1-deficient phagocytes showed elevated *Mtb* infection rate and increased *Mtb* growth in comparison to WT cells resulting in increased mycobacterial numbers *in vitro* after 4 days post-infection. These observations are accompanied by findings that demonstrate that IRG1-deficient macrophages and dendritic cells (DCs) have increased levels of lipid droplets (LDs), which are reservoirs of host nutrients for *Mtb*. Therefore, our findings demonstrate that IRG1 is a major player in controlling the acute phase of *Mtb* infection by regulating inflammatory responses and availability of host nutrients.

## Results

We explored the physiological relevance of IRG1 deficiency during *Mtb* infection *in vivo* using an infection model established in C57Bl/6 mice. A previous study demonstrated that, compared to WT mice, IRG1-deficient mice (IRG1^-/-^) were more susceptible to aerosol infection with the *Mtb* Erdman strain [32]. Here, we comparatively evaluated the impact of an intranasal inoculation of WT and IRG1^-/-^ mice by a high dose (10^5^ CFU per mouse) of the virulent *Mtb* H37Rv strain, the attenuated strain *Mtb* H37Ra and the vaccinal mycobacterial strain *M. bovis* BCG 1173P2 (**Figure 1A**). We monitored pathologic parameters, changes in body weight and survival as well as mycobacterial burden during the course of infection for 84 days post-infection (dpi). During the first two weeks post-infection, no apparent differences in body weight of WT and IRG1^-/-^ mice, infected with the different mycobacterial strains, were apparent (**Figure 1B**). From 14 dpi onwards, IRG1^-/-^ mice exposed to *Mtb* H37Rv started to rapidly lose weight (approximately one gram daily), while WT mice exposed to virulent and attenuated strains, as well as IRG1*^-/-^* mice exposed to *Mtb* H37Ra and BCG were not affected (**Figure 1B**). The rapid loss in body weight of *Mtb* H37Rv-infected IRG1*^-/-^* mice was accompanied by other clinical symptoms, such as shortness of breath and lethargy, and the mice reached moribund conditions and died between 3 to 4 weeks post-infection (**Figure 1C**), similar to IRG1^-/-^ mice infected by *Mtb* Erdman [32]. In contrast, all infected WT mice as well as IRG1^-/-^ mice exposed to *Mtb* H37Ra and BCG did not display loss in body weight nor morbidity and mortality, which we followed up until 84 dpi (**Figure 1B-C**). These observations confirm a substantial susceptibility of mice to virulent *Mtb* infection in the absence of functional IRG1. Therefore, we next focussed on immunopathology and mycobacterial burden of infected organs of the differently infected WT and IRG1^-/-^ mice.

**Figure 1.**
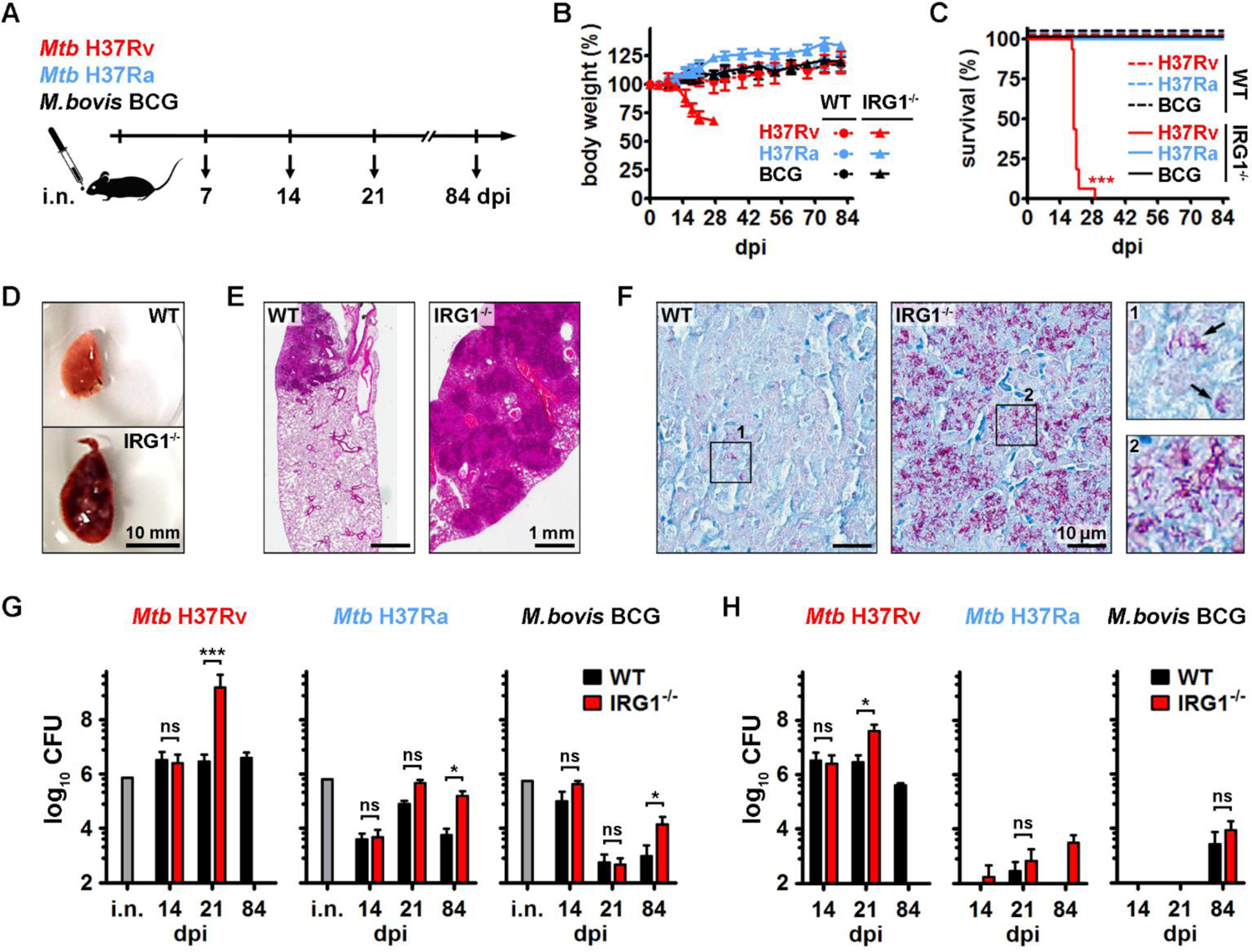
IRG1-deficient mice are highly susceptible to infection by virulent *Mtb*. **(A)** Workflow of the experimental setup during intranasal (i.n.) infection of WT mice and IRG1^-/-^ mice with the indicated mycobacterial strains and the different read-out time points at days post-infection (dpi). **(B)** Changes in body weight of WT mice and IRG1^-/-^ mice and **(C)** their survival rates after intranasal administration of *Mtb* H37Rv (red), *Mtb* H37Ra (blue) and *M. bovis* BCG (black). **(D)** Photographs of representative examples of the left lung lobe of WT mice and IRG1^-/-^ mice infected by *Mtb* H37Rv at 21 dpi. **(E)** Immunopathology of mouse lungs infected by *Mtb* H37Rv at 21 dpi, as determined by hematoxylin phloxine saffron staining of histological sections. **(F)** The mycobacterial load in lungs of WT mice and IRG1^-/-^ mice infected by *Mtb* H37Rv at 21 dpi was determined by acid-fast stain (Ziehl-Neelsen staining) of representative histological sections. Insets display labeled mycobacteria in purple (arrows). **(G)** Determination of the mycobacterial burden of the left lung lobe of WT and IRG1^-/-^ mice by counting colony-forming units (CFU) of administered intranasal inoculums (i.n.) and the indicated mycobacterial strains at different days post-infection. **(H)** CFU determination of spleens of mice infected by the indicated mycobacterial strains. Shown are mean ± SD. Statistical differences were determined by survival Log-rank (Mantel-Cox) test (C) and by non-parametric Mann-Whitney test (G, H). ns: non-significant, * P value < 0.05, ** P value < 0.01, *** P value < 0.001.

At 21 dpi, the lungs of *Mtb* H37Rv-infected IRG1^-/-^ mice were enlarged and displayed macroscopic pathological features with areas of necrotizing lesions (**Figure 1D**), while these observations were absent in lungs of mice infected with *Mtb* H37Ra or BCG (**Figure S1**). Histological examination of the lungs using hematoxylin phloxine saffron staining showed massive granulomatous inflammation in *Mtb* H37Rv-exposed IRG1^-/-^ mice compared to their WT counterparts (**Figure 1E**). Ziehl-Neelsen staining of mycobacteria applied to the histological lung sections detected substantial amounts of invading *Mtb* H37Rv in the lung parenchyma of IRG1^-/-^ mice (**Figure 1F, inset 2**), while lungs of WT mice contained much lower numbers of bacteria (**Figure 1F, inset 1**). Determination of pulmonary mycobacterial loads at different time points post-*Mtb* H37Rv-infection demonstrated that, while the load is comparable after two weeks of infection (14 dpi), a 3-Log increase in CFU were detected in the lungs of IRG1^-/-^ mice compared to WT mice at 21 dpi (**Figure 1G**). After infection with a similar inoculum of *Mtb* H37Ra or BCG, IRG1^-/-^ mice are not more susceptible for the first 3 weeks. Nevertheless, an increase in the bacterial load in deficient mice is observed during the chronic phase of infection (84 dpi) without any clinical manifestations (**Figure 1G**). In addition, examination of spleen showed *Mtb* H37Rv susceptibility in IRG1^-/-^ mice 21 dpi (**Figure 1H**). A lower level of bacteria was detected in the spleen of BCG and *Mtb* H37Ra infected mice showing the difficulty of these attenuated strains to disseminate in peripheral organs. In the spleen of *Mtb* H37Ra infected mice, bacteria are only detected in IRG1^-/-^ mice 84 dpi (**Figure 1H**).

Altogether, these results indicate a major contribution of IRG1 in the antimicrobial host response not only at the mucosal site of infection but also systemically. Our observations in vivo clearly indicate that IRG1^-/-^ mice are highly susceptible to virulent *Mtb* developing a severe phenotype leading to the death of all animals within 3-4 weeks. Infection of IRG1^-/-^ mice with an attenuated mycobacterial strain (*M. bovis* BCG or *Mtb* H37Ra) leads to lower pathogenicity, but these mice still show a higher susceptibility compared to infected WT mice at a late stage post infection. We performed immune cell characterization in WT and IRG1^-/-^ mice during *Mtb* H37Rv infection until 21 dpi by determining the total cell numbers of macrophages, eosinophils, DCs, neutrophils, CD4 T cells, CD8 T cells and B cells in lungs (**Figure S2A**) and spleen (**Figure S2B**). Until 14 dpi, all lung and splenic cell populations were comparable between WT and IRG1^-/-^ mice and did not show remarkable differences (**Figure 2A, Figure S2C**). In contrast, the time period between 14 dpi and 21 dpi led to a dramatic increase in pulmonary macrophages (both alveolar and interstitial ones) as well as neutrophils in the lungs of IRG1^-/-^ mice, while levels of CD4 T cells, CD8 T cells and B cells were significantly decreased compared to WT mice (**Figure 2A**). In the spleen, DCs and neutrophils were significantly increased in IRG1^-/-^ mice at 21 dpi, while CD4 T cells and B cells exhibited a reduction compared to WT cells (**Figure S2C**). These findings demonstrate that macrophages (in the lung), DCs (in the spleen) and neutrophils (in both organs) are the main cell populations that are responsible for the increased progression of *Mtb* infection in IRG1^-/-^ mice. This is underlined by the fact that neutrophils in the lungs of IRG1^-/-^ mice displayed a dramatically higher mycobacterial burden compared to lungs of their WT counterparts, when they were labelled in situ (**Figure 2B**). In line with this, the observed reduction in T cell and B cell levels in both, lung and spleen, of IRG1^-/-^ mice might suggest impaired adaptive immunity in those animals compared to *Mtb*-infected WT mice, which could be beneficial to *Mtb* progression and immunopathology observed in IRG1^-/-^ animals.

**Figure 2.**
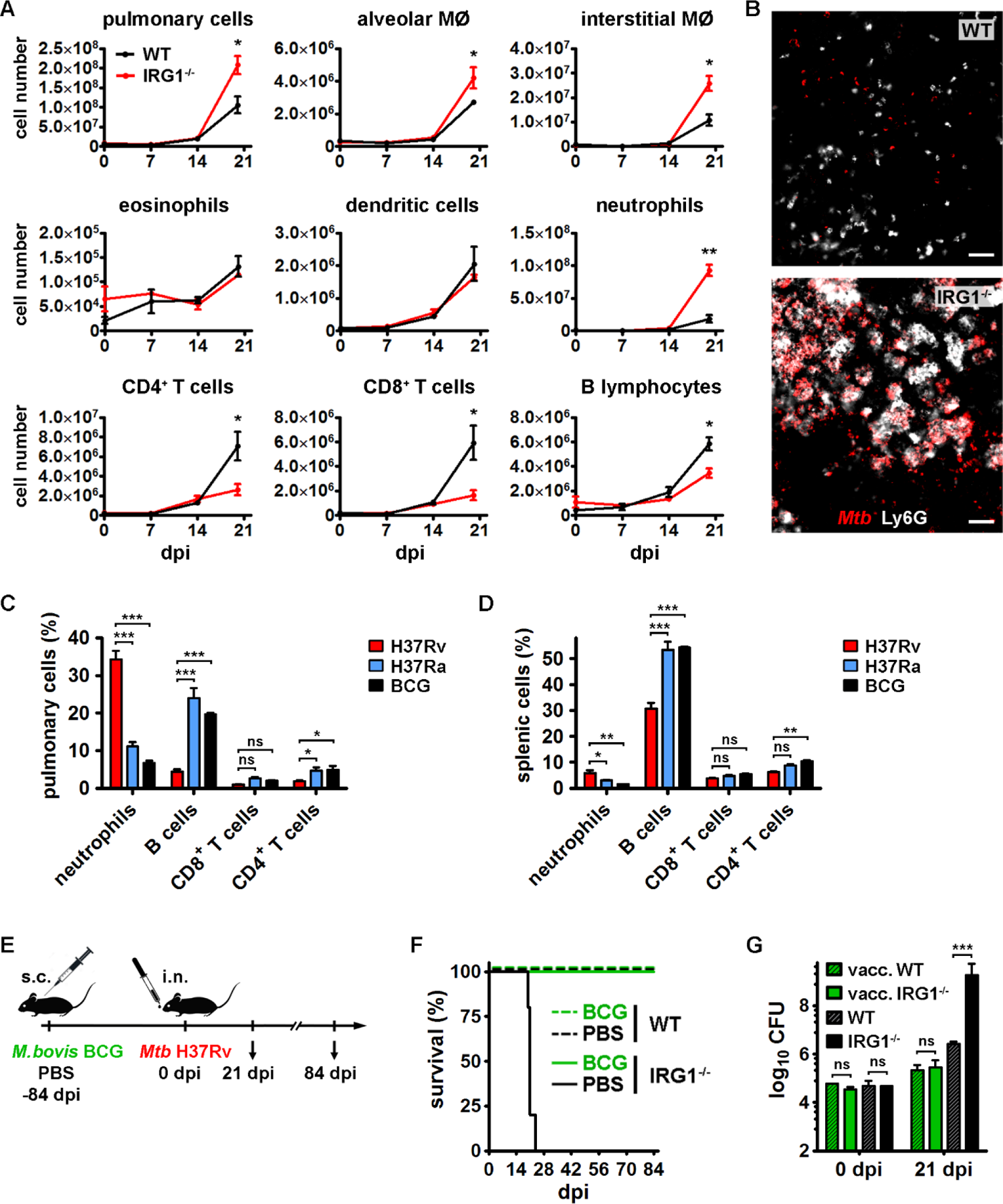
Adaptive immune responses help to overcome susceptibility of IRG1^-/-^ mice to virulent *Mtb* infection. **(A)** Profiling of lung immune cells of WT mice and IRG1^-/-^ mice during the course of *Mtb* H37Rv infection. Histograms depict changes in cell numbers of the indicated immune cell populations, as determined by flow cytometry using specific cell surface markers. All cell numbers were normalized to the total cell number analyzed in each sample and were extrapolated to the whole organ. Shown are mean ± SEM of cells obtained from three individual mice per group. **(B)** Representative images of lung tissue cryosections obtained from *Mtb* H37Rv-infected WT mice and IRG1^-/-^ mice at 21 dpi and acquired by confocal microscopy. Mycobacteria (red) were labeled with a specific lipoarabinomannan (LAM) antibody, while neutrophils (white) were labeled with an antibody against Ly6G. Bar: 50 µm. **(C-D)** Differences in neutrophil numbers and adaptive immune cell populations of IRG1^-/-^ mice infected by *Mtb* H37Rv (red), *Mtb* H37Ra (blue) and *M. bovis* BCG (black). Shown are percent levels of the indicated cell types in **(C)** pulmonary cell populations and **(D)** splenic cell populations at 21 dpi. **(E)** Workflow of the experimental setup of vaccination of WT mice and IRG1^-/-^ mice by subcutaneous (s.c.) injection of *M. bovis* BCG followed by intranasal (i.n.) challenge with *Mtb* H37Rv. Control mice received s.c. injections of PBS. **(F)** Survival rate of BCG-vaccinated mice (green) and PBS-injected control mice (black) after intranasal challenge with *Mtb* H37Rv at a similar dose shown in Figure 1C. **(G)** Determination of mycobacterial loads in left lung lobes of BCG-vaccinated (vacc.) and control WT and IRG1^-/-^ mice by CFU counting 21 days after challenge with *Mtb* H37Rv. Shown are mean ± SD. Statistical differences were determined by non-parametric Mann-Whitney test (A, G), by ANOVA and Bonferroni post-hoc correction for multiple comparisons (C, D) and survival Log-rank (Mantel-Cox) test (F). ns: non-significant, * P value < 0.05, ** P value < 0.01, *** P value < 0.001.

Immune cell profiling was also characterized during infection of IRG1^-/-^ mice with attenuated *Mtb* H37Ra and BCG. At 21 dpi, the number of recruited neutrophils is much lower in both lungs (**Figure 2C**) and spleen (**Figure 2D**) for mice infected with the attenuated strains compared to mice infected with *Mtb* H37Rv. While infection with the virulent strain induces a decrease in the number of lymphocytes, it was observed that these cells were more prevalent in lungs (**Figure 2C**) and the spleen (**Figure 2D**) of mice infected with the attenuated strains. Together, our observations indicate that the high susceptibility of IRG1^-/-^ mice infected with virulent *Mtb* is associated with high tissue neutrophils count and a failure to recruit adaptive immune response cells.

Since the T and B cell response appears to be diminished in infected IRG1^-/-^ mice, we investigated whether these mice had difficulties to establish a protection after vaccination. To do this, C57BL/6 WT and IRG1^-/-^ mice were subcutaneously vaccinated with the *M. bovis* BCG Pasteur vaccine 1173P2. Three months later, the mice were challenged with an intranasal dose of *Mtb* H37Rv similar to previous infection experiments to monitor their degree of protection compared to unvaccinated mice (**Figure 2E**). We first observed that *Mtb* infection was no longer lethal in vaccinated IRG1^-/-^ mice contrary to non-vaccinated ones (**Figure 2F**). These findings imply that IRG1^-/-^ mice are able to develop, like WT mice, a protective immune response after BCG vaccination. To confirm this, we evaluated the pulmonary bacterial load of *Mtb* 21 dpi in WT and IRG1^-/-^ mice, pre-vaccinated or not. The results show that, while unvaccinated IRG1^-/-^ mice are much more susceptible than WT mice, both strains of mice show a similar level of protection in the lungs after vaccination with BCG (**Figure 2G**). Taken together, this means that, while IRG1 plays a very important role in the control of primary infection, it does not impair vaccine protection establishment. Given the critical role of CD4^+^ and CD8^+^ T cells in protection against *Mtb*, it was rather surprising to discover that BCG-vaccinated, IRG1^-/-^ mice were as resistant WT mice to an *Mtb* challenge. This suggests that IRG1 may also affect phagocytes, which are important mediators for the generation of protective innate immunity against TB [33] and led us to further investigate the cell biology of IRG1^-/-^ phagocytes.

Given the role of IRG1 as a mitochondrial enzyme of the TCA cycle that is induced upon inflammation and infection, we first studied the metabolic activity of bone marrow-derived macrophages (BMDMs) derived from WT and IRG1^-/-^ mice by microscale oxygraphy using the Seahorse approach. To probe the glycolytic activity of resting and activated BMDMs, glucose, oligomycin A, and 2-deoxyglucose (2-DG) were sequentially injected and the extracellular acidification rate (ECAR) was measured (**Figure 3A**). While injected glucose is serving to feed glycolysis, oligomycin A is inhibiting ATP synthase in the electron transport chain and changes the ATP/ADP ratio [34]. The addition of 2-DG is inhibiting glycolysis and terminates the experiment and therefore provides a non-glycolytic measurement of ECAR at baseline levels. We observed a high glycolytic reserve capacity in resting IRG1^-/-^ macrophages compared to resting WT cells demonstrated by increased ECAR values over the entire measurement period (**Figure 3A**). Also, upon activation of BMDMs by overnight stimulation with LPS and IFNγ, IRG1^-/-^ cells exhibited elevated ECAR values compared to their WT counterparts. To test mitochondrial respiration of cells as an indicator of oxidative phosphorylation activity, we also performed measurements of the oxygen consumption rate (OCR). Injection of oligomycin A decreases electron flow and leads to a reduction of mitochondrial respiration, which is followed by addition of carbonyl cyanide-4 (trifluoromethoxy) phenylhydrazone (FCCP), a protonophoric uncoupling agent that is collapsing the proton gradient and is disrupting the mitochondrial membrane potential [35]. The terminal injection of rotenone and antimycin A shuts down mitochondrial respiration and allows the calculation of non-mitochondrial OCR. IRG1^-/-^ macrophages did not show differences (or only very minor, negligible ones) in their OCR compared to WT cells at both resting and activated conditions (**Figure 3B**). In addition, we also calculated the mitochondrial spare respiratory capacity (SRC), which is the difference between maximal and basal respiration, that correlates with the bio-energetic adaptability of mitochondria in response to pathophysiological stress conditions [36]. We found that IRG1^-/-^ macrophages have an increased SRC at both resting and activated conditions compared to WT macrophages (**Figure 3C**). Together, these data show that IRG1^-/-^ macrophages are characterized by a more glycolytic metabolism, and that their mitochondria remain highly dynamic to meet extra energy requirements (e.g. in response to acute cellular stress and/or infection by pathogens), as displayed by their increased SRC levels.

**Figure 3.**
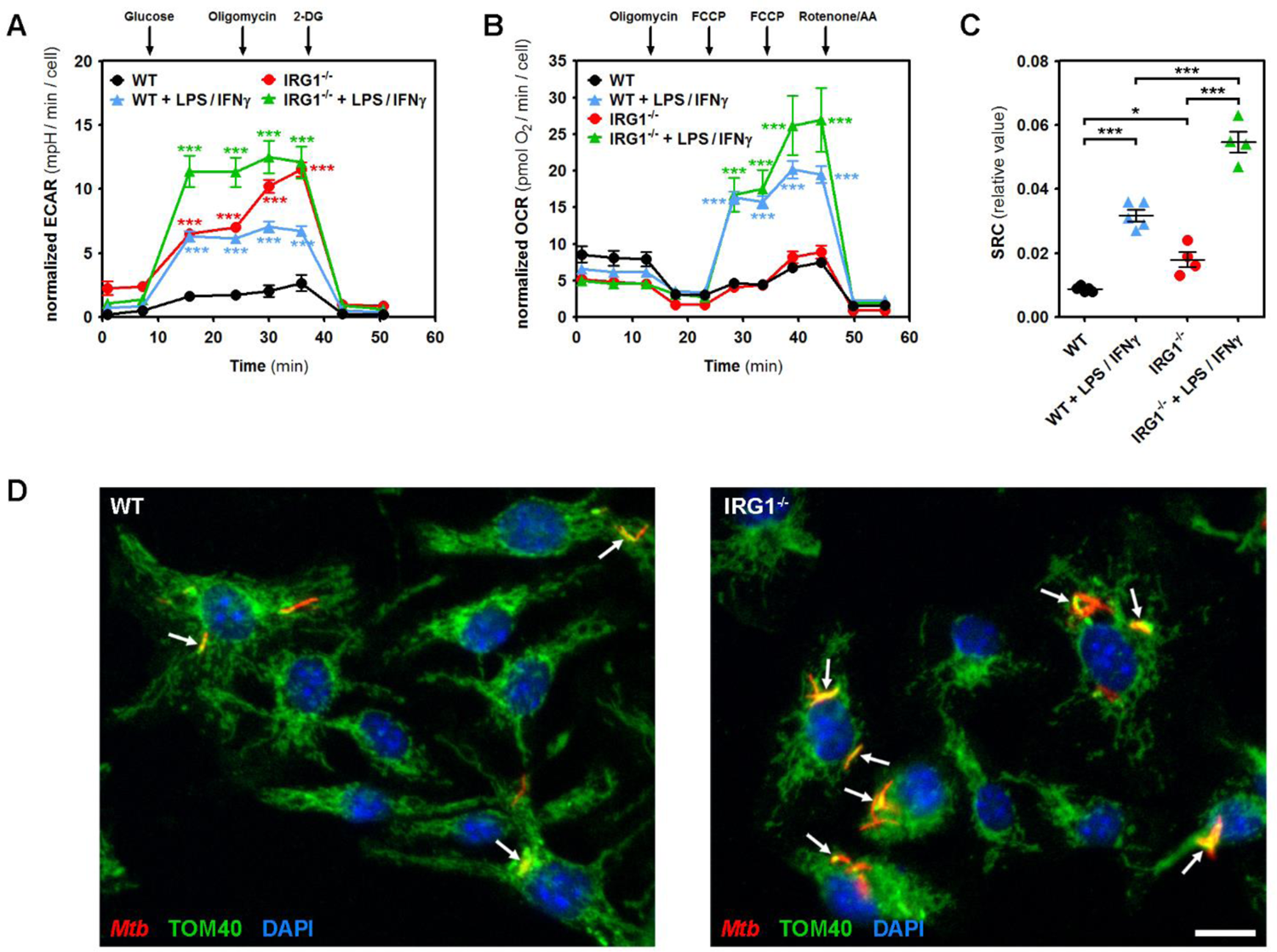
IRG1^-/-^ macrophages are characterized by a more glycolytic metabolism, and *Mtb* are found in close vicinity of mitochondria of both, WT and IRG1^-/-^ cells. (A-C) Glycolysis and mitochondrial respiration were measured by Seahorse analysis determining **(A)** extracellular acidification rate (ECAR) and **(B)** oxygen consumption rate (OCR) of bone marrow-derived macrophages (BMDM), respectively. Cells were analyzed in resting conditions and after stimulation with 100 ng/ml LPS and 20 ng/ml IFNγ for 18 h. Injection of substrates and different inhibitors occurred at the indicated time points (more details in the methods section). **(C)** Bio-energetic plasticity of mitochondria of BMDMs in response to pathophysiological stress conditions was calculated by their spare respiratory capacity (SRC). 2-DG: 2-deoxy-D-glucose, FCCP: carbonyl cyanide-4 (trifluoromethoxy) phenylhydrazone, AA: antimycin A. **(D)** BMDMs obtained from WT mice (left) and IRG1*^-/-^* mice (right) were infected by Mtb H37Rv-DsRed (red), fixed at 48 hpi and analyzed by confocal microscopy. Mitochondria were labeled with an antibody against TOM40 (green), a marker of the mitochondrial outer membrane, and nuclei were stained with DAPI (blue). Arrows indicate *Mtb* in close vicinity to mitochondria. Bar: 10 µm. Statistical differences between matching WT and IRG1*^-/-^* samples were determined by two-way ANOVA with a 95% confidence interval followed by Bonferroni’s post-test (A-C). * P value < 0.05, *** P value < 0.001.

Next, we analyzed the subcellular distribution of mitochondria in *Mtb*-infected BMDMs by confocal microscopy using a marker against TOM40, an import channel of the mitochondrial outer membrane. Mitochondria in both, WT and IRG1^-/-^ macrophages, were found distributed over the entire cell periphery (**Figure 3D**) with mitochondria localized in close vicinity of intracellular *Mtb* (arrows in **Figure 3D**), as reported previously (summarized in [37]). We noted more bacteria in infected IRG1^-/-^ macrophages compared to WT cells, but found mitochondria surrounding *Mtb*-containing vacuoles in both cell types. These findings suggest that IRG1 deficiency does not directly affect mitochondria morphology and possible interactions between *Mtb* and mitochondria.

To further examine the contribution of IRG1 during *Mtb* infection *in vitro*, we infected WT and IRG1^-/-^ BMDMs by the virulent *Mtb* H37Rv strain and investigated the profile of *Irg1* gene expression by quantitative RT-PCR during the course of infection. While *Irg1* was not expressed in resting, non-infected BMDMs, and, as expected, in IRG1^-/-^ macrophages, the stimulation of WT cells, but not of IRG1^-/-^ cells, with 100 ng/ml LPS and 20 ng/ml IFNγ for 24 h induced *Irg1* expression considerably (**Figure 4A**), in accordance with previous observations [19]. Infection by *Mtb* H37Rv (at MOI=1) also rapidly induced *Irg1* expression in WT BMDMs and showed increased levels between 2 h and 48 h post-infection (hpi), with almost undetectable levels at 96 hpi, comparable to non-infected cells (**Figure 4A**). Full absence of *Irg1* signals in BMDMs obtained from IRG1^-/-^ mice showed the high specificity of the RT-PCR approach (**Figure 4A**). In accordance with our gene expression data, we detected the IRG1 protein by western blotting in lysates of *Mtb* H37Rv-infected WT BMDMs at 24 hpi, 48 hpi and at lower levels at 72 hpi (**Figure 4B**). In addition to BMDMs, LPS stimulation also induced IRG1 expression in bone marrow-derived dendritic cells (BMDCs), as detected in total cell lysates (TCL) from resting *versus* LPS-stimulated BMDCs (**Figure 4C**).

**Figure 4.**
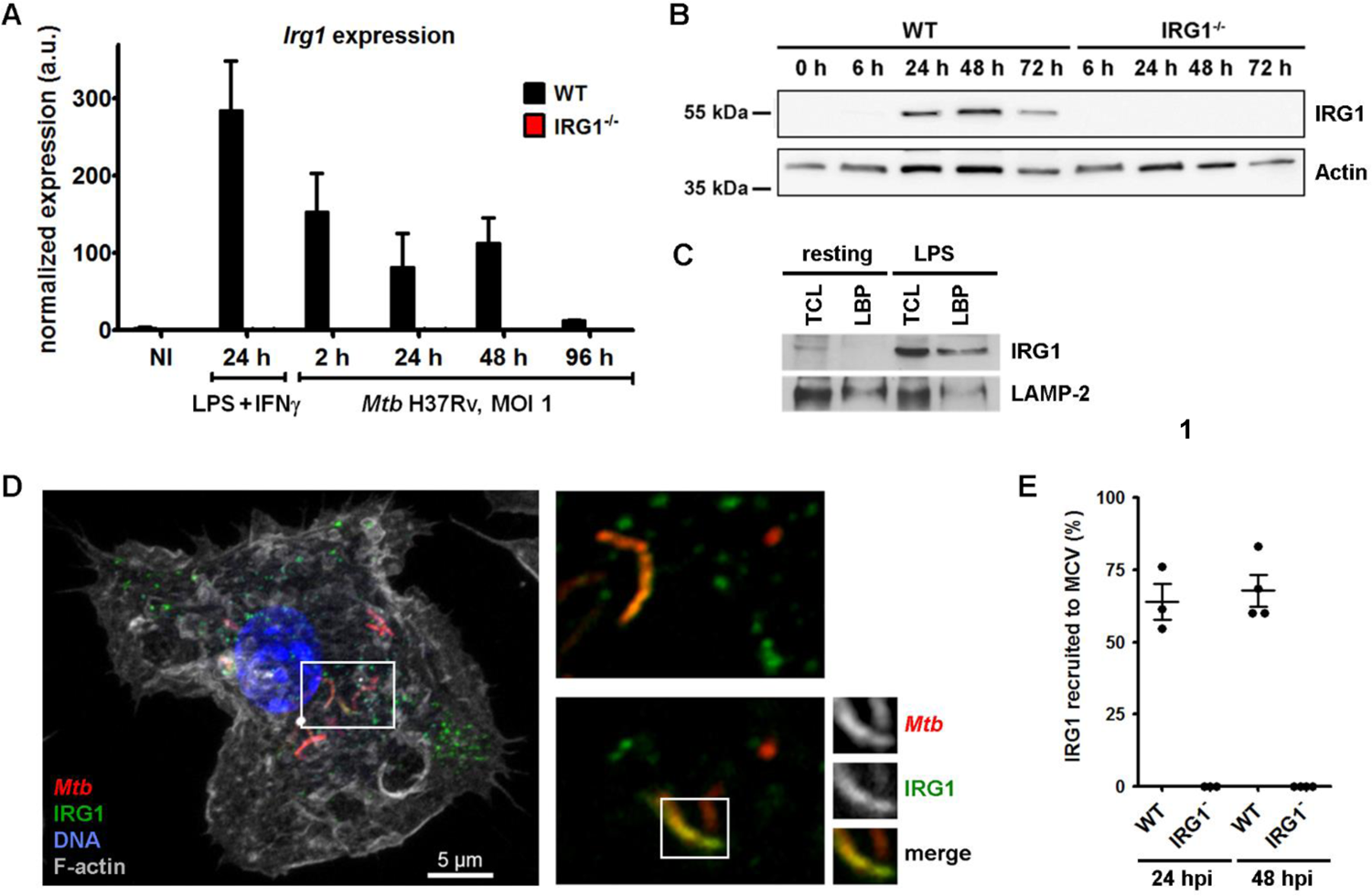
*Irg1* expression is induced by *Mtb* infection and IRG1 is recruited to *Mtb*-containing vacuoles. **(A)** BMDMs of WT mice and IRG1^-/-^ mice were either treated with 100 ng/ml LPS and 20 ng/ml IFNγ or infected with *Mtb* H37Rv for the indicated time points. Non-infected (NI) cells served as control. Transcription of *Irg1* was assessed by quantitative RT-PCR and normalized to the expression of glyceraldehyde-3-phosphate dehydrogenase *(Gapdh)*. Shown are mean ± SEM obtained from BMDMs of three different mice. **(B)** Expression of IRG1 protein in total cell lysates (TCL) of WT and IRG1^-/-^ BMDMs infected with *Mtb* H37Rv for the indicated time points as detected by western blotting. Expression of Actin is shown as loading control. One experiment representative of three independent ones is shown. **(C)** Detection of IRG1 and LAMP-1 by western blotting in TCL and purified fractions of latex bead-containing phagosomes (LBP) of bone marrow-derived dendritic cells (BMDC) obtained from WT mice. BMDCs were analyzed at resting state (left) or 16 h after the addition of 100 ng/ml LPS (right). **(D)** Confocal image of the subcellular localization of *Mtb* H37Rv-DsRed (red) and IRG1 (green) in WT BMDM at 24 h post-infection (hpi). The nucleus was labeled with DAPI (blue), while F-actin was visualized by Phalloidin staining (gray). Insets (right panel) depict different focal planes of the same area showing co-localization of IRG1 to *Mtb* phagosomes. **(E)** Recruitment of IRG1 to *Mtb*-containing vacuoles (MCV) in WT BMDMs and IRG1^-/-^ BMDMs was assessed by confocal microscopy and by manual analysis of focal planes of entire cells. Shown are mean ± SEM of 3 (24 hpi) and 4 (48 hpi) independent experiments analyzing at least 30 infected cells per condition.

Intracellular pathogens, such as *Mtb*, that enter immune cells by phagocytosis, are located in phagosomes, which further interact with endosomal compartments during phagosome maturation. In a previous study, using a well-established phagocytosis model system of antigen-coupled beads, we identified by quantitative mass spectrometry the specific recruitment of IRG1 to phagosomes of LPS-stimulated BMDCs [38]. We confirmed this observation by western blotting analyzing purified latex bead-containing phagosomes (LBP). We found IRG1 present in LBP lysates of LPS-stimulated BMDCs, while it was absent in phagosomal lysates of resting BMDCs (**Figure 4C**). In contrast, lysosome-associated membrane protein 1 (LAMP-1), a membrane glycoprotein originated from late endosomes and lysosomes, was recruited to LBPs of both, resting and LPS-stimulated BMDCs (**Figure 4C**). In addition, previous work on macrophage infection by *Legionella pneumophila* showed that IRG1 could also be recruited to *Legionella*-containing phagosomes [23]. Therefore, we examined a possible association of IRG1 with *Mtb*-containing vacuoles (MCVs). *Mtb*-infected BMDMs were fixed 24 hpi and labeled for IRG1, F-actin and nuclei for analysis by confocal microscopy (**Figure 4D**). IRG1 was found recruited to MCVs, as several IRG1 signals (green) co-localized with *Mtb* signals (red) (**insets of Figure 4D**). In other parts of the cell, IRG1 remained located outside of those vacuoles and was restricted to other cell organelles, such as mitochondria, as demonstrated previously during pro-inflammatory conditions [38]. We further quantified the recruitment of IRG1 to MCVs by confocal microscopy at 24 hpi and 48 hpi, which was evident in 50% to 80% of all analyzed MCVs of WT cells dependent on the investigated time point and experiment, while no signal was found at MCVs of IRG1^-/-^ macrophages (**Figure 4E**). These findings demonstrated that IRG1 is induced upon *Mtb* infection, followed by its recruitment to *Mtb*-containing phagosomes, suggesting a role of IRG1 in the intracellular host defense against mycobacteria.

To further elucidate this role in murine phagocytes, we infected BMDMs and BMDCs derived from WT mice or IRG1^-/-^ mice with a GFP-expressing *Mtb* H37Rv strain and followed colonization of host cells and *Mtb* replication up to 96 hpi (**Figure 5A**). Cells were grown in 384-well plates and their nuclei were labeled to enable analysis by an automated confocal microscopy approach using in-house multiparametric imaging that allowed acquisition and examination of hundreds of images generating robust and reproducible data sets (**Figure S3A**) [40]. Algorithms were applied to input images, which resulted in segmentation of the different fluorescent signals allowing nuclei detection, cell and bacteria selection and further downstream analysis to determine infection rate and number of bacteria per cell (**Figure S3A**). First, we observed that the efficacy of *Mtb* uptake by WT and IRG1^-/-^ phagocytes were comparable, as determined by the percentages of *Mtb*-infected BMDMs and BMDCs at 2 hpi (**Figure 5B**). We then compared the *Mtb* intracellular area per infected cell, which directly correlates with the number of *Mtb* per cell, which showed no differences between WT and IRG1^-/-^ BMDMs and BMDCs at 2 hpi (**Figure 5C**). In contrast, at 96 hpi the percentages of *Mtb*-infected BMDMs and BMDCs were largely increased in IRG1^-/-^ BMDMs and BMDCs (**Figure 5B**), showing that IRG1 deficiency favors survival and/or intracellular growth of *Mtb*. This was further supported by the fact that at 96 hpi IRG1-deficient BMDMs and BMDCs also had significantly higher *Mtb* numbers per cell compared to WT phagocytes (**Figure 5C**). These findings indicate that the expression of IRG1, induced by *Mtb* infection, and the presence of IRG1 during the course of infection enable host cells to restrict excessive growth and replication of *Mtb*.

**Figure 5.**
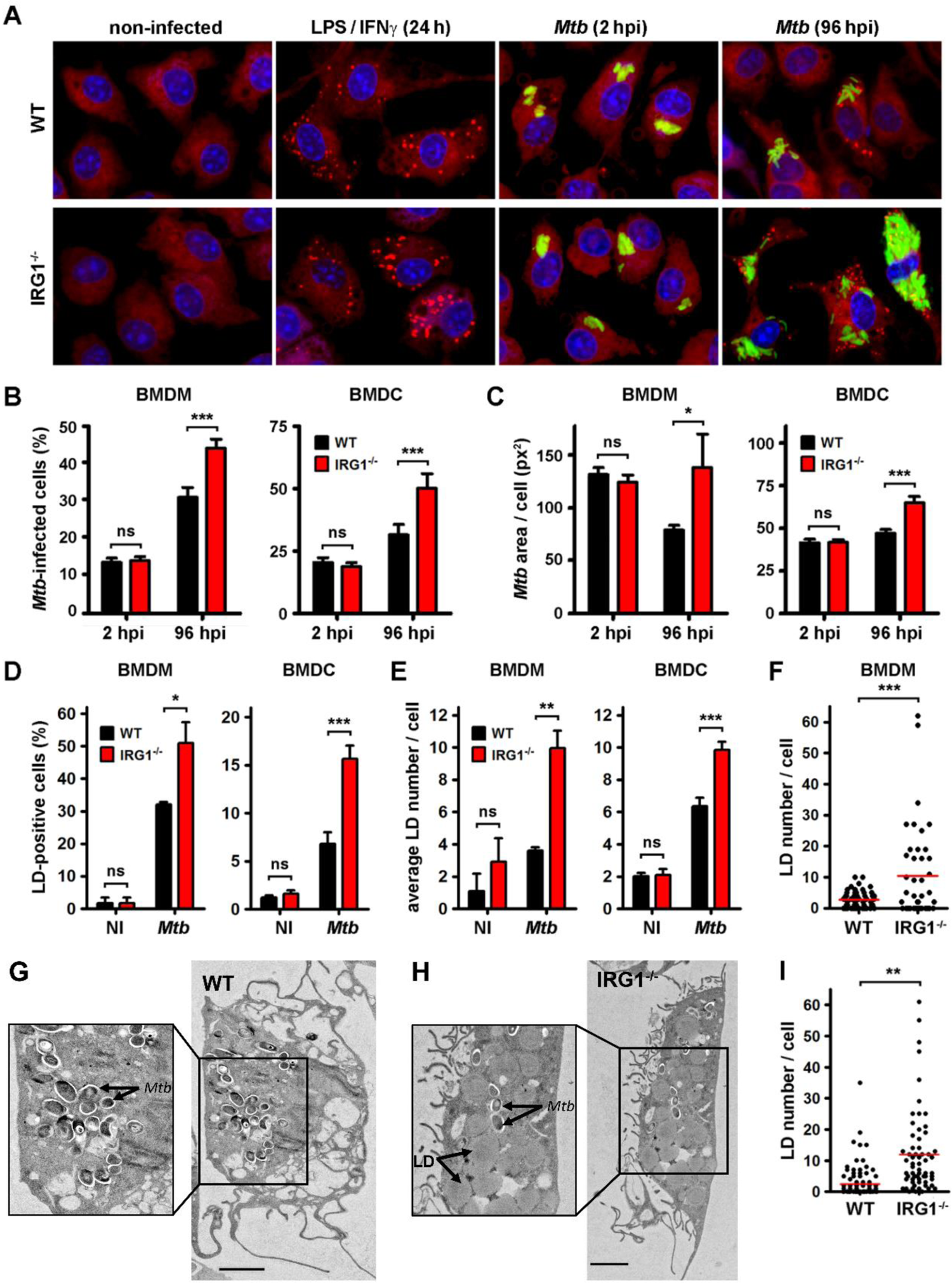
IRG1 deficiency leads to increased *Mtb* replication and increased amounts of lipid droplets (LDs) in phagocytes. **(A)** WT BMDMs (upper panel) and IRG1^-/-^ BMDMs (lower panel) were left non-infected (NI) or infected with *Mtb* H37Rv-GFP (green) for the indicated time points. Alternatively, cells were also treated with 100 ng/ml LPS and 20 ng/ml IFNγ for 24 h. All cells were fixed and labeled with DAPI and LipidTOX DeepRed to stain nuclei (blue) and LDs (red), respectively. Shown are representative images acquired by automated confocal microscopy and image analysis. Bar: 10 µm. **(B-F)** BMDMs and BMDCs were grown in 384-well plates, infected with *Mtb* H37Rv-GFP (green) and analyzed by automated confocal microscopy. **(B)** Histograms displaying the percentage of infected BMDMs (left panel) and BMDCs (right panel) at 2 hpi and 96 hpi. **(C)** Histograms showing the bacterial area per infected BMDM (left panel) and infected BMDC (right panel) at both time points, which directly correlates with the number of *Mtb* per infected cell. **(D)** Histograms depicting the percentage of LD-positive NI and *Mtb*-infected BMDMs (left panel) and BMDCs (right panel) at 96 hpi, as determined by automated confocal microscopy. **(E)** Histograms showing the average LD number per cell for both cell types at 96 hpi. Shown are mean ± SEM of at least 6 analyzed wells per condition (n > 400 cells) of one representative out of three independent experiments. **(F)** LD numbers of BMDMs at 96 hpi (n=100) were also counted manually in a blinded fashion to verify accuracy of the automated image analysis pipeline. **(G-I)** Ultrastructural analysis of LD formation in *Mtb*-infected BMDMs derived from WT mice **(G)** and IRG1^-/-^ mice **(H)** at 96 dpi by electron microscopy. Bar: 2 µm. **(I)** Numbers of lipid droplets (indicated as LD in the inset) were quantified in WT cells (n=100) and IRG1^-/-^ cells (n=64). Statistical differences were determined by Student’s t-test (ns: non-significant, * P value < 0.05, ** P value < 0.01, *** P value < 0.001).

Previous observations have shown that *Mtb* persistence/replication relies on the availability of host nutrients that the bacteria exploit to build their replicative niche in phagocytes. During infection, the key carbon source for intracellular *Mtb* consists of host lipids that are mainly stored in LDs and are responsible of the foamy phenotype of macrophages inside tuberculosis granuloma [41]. Therefore, we investigated the impact of IRG1 deficiency on the formation of LDs and their availability in host cells during *Mtb* infection *in vitro*. We used a specific dye to label neutral lipids, predominantly located in LDs, in BMDMs and BMDCs at 96 hpi, i.e. the time point where differences in *Mtb* replication rates were very prominent. We again applied automated confocal microscopy and multiparametric image analysis to detect and quantify LDs (**Figure S3B**), which were visualized in both cell types (**Figure 5A**). While non-infected BMDMs and BMDCs exhibited very low percentages of LD-positive cells (**Figure 5D**) or low LD numbers per cell (**Figure 5E**), *Mtb* infection increased the rate of LD-positive cells and LD numbers per cell. The percentage of LD-positive cells was further increased in the IRG1^-/-^ BMDMs and BMDCs compared to WT cells (**Figure 5D**), and those cells showed significantly more LDs per cell (**Figure 5E**). While infected WT BMDMs displayed on average 2.7 ± 0.3 LDs per cell, infected IRG1^-/-^ BMDMs contained on average 10.4 ± 2.3 LDs per cell (**Figure 5F**).

Strikingly, bystander non-infected cell also had increased lipid droplets numbers (**Figure S4**). These results demonstrate clearly that the presence of IRG1 is concomitant with the reduction of LDs in macrophages and DCs during *Mtb* infection. Finally, we also analyzed LD formation in infected BMDMs on the ultrastructural level by performing electron microscopy (**Figure 5G-H**). Similar to our findings by confocal microscopy, we found significantly more LDs per cell in IRG1^-/-^ BMDMs compared to WT cells (**Figure 5I**). Altogether, our data show a correlation of LD formation upon infection and the impact on *Mtb* replication in phagocytes *in vitro*, which is favored in conditions where IRG1 is absent.

## Discussion

In this study, we addressed the role of IRG1 in immunometabolic host responses during mycobacterial infections and were able to demonstrate that expression of IRG1 is essential to dampen immunopathology, both *in vitro* and *in vivo*. Our results on IRG1-deficient mice show that upon infection by a virulent *Mtb* strain (H37Rv), but importantly not by attenuated *Mtb* (H37Ra) and by the vaccinal strain *M. bovis* BCG, infected animals display high susceptibility and mortality with exacerbated *Mtb* H37Rv loads in lung and spleen at 3-4 weeks post-infection. Moreover, the immunopathological features of H37Rv-infected IRG1^-/-^ mice (increased inflammation and neutrophil infiltration in lungs and spleens) are similar to those reported previously by the Stallings lab [32]. In that study, the authors investigated the impact of IRG1 on *Mtb* infection using a different virulent *Mtb* strain (Erdman) and a different route of inoculation (aerosol versus intranasal infection like in our study). IRG1 expression is also essential to control other virulent bacterial or viral infections, as it was shown previously for *Legionella pneumophila* [23, 24], *Salmonella enterica* [28] as well as for *Brucella melitensis* and *B. abortus* [30, 31]. While IRG1^-/-^ mice were more susceptible to *Brucella* infection and rescued the virulence defect of a *S. enterica* mutant, infection of those mice by *Listeria monocytogenes* and influenza A virus did not result in altered susceptibility of infected IRG1^-/-^ animals [32]. In our study, we provide evidence that IRG1^-/-^ mice are only highly susceptible to virulent *Mtb* but not to attenuated mycobacterial species, such as the vaccinal strain *M. bovis* BCG or the attenuated *Mtb* strain H37Ra. The difference in susceptibility depending on microbes and strain pathogenicity suggests the contribution of microbial effectors in the increased pathogenicity linked to the impairment of IRG1 functions. In the context of *Mtb*, the main difference between the two *Mtb* strains (H37Ra and H37Rv) is linked to the secretion of important effectors of the ESX-1 type VII secretion system (T7SS) through the regulation by the transcriptional regulator PhoP [42]. Future studies would need to clarify the relevance of the ESX-1 T7SS and its virulence effectors on altered IRG1 functions by comparing WT and IRG1-deficient mice.

Profiling of the different pulmonary immune cell populations after 21 dpi of *Mtb* H37Rv infection, which is corresponding with the end of the acute phase of infection, revealed a massive infiltration of neutrophils, which was accompanied by increased numbers of alveolar and interstitial macrophages. In TB, neutrophils have deleterious as well as host-beneficial roles, and when they are overloaded with bacilli, their abundance in infected tissues correlates with disease severity [43]. In lungs and spleens of IRG1^-/-^ mice, we also observed reduced amounts of CD4^+^ and CD8^+^ T cells, which has not been reported by Nair and colleagues [32], and highlights a possible impact of IRG1 deficiency on the onset of adaptive immunity. Recent findings support the regulatory role of the Irg1/itaconate pathway in adaptive immune responses during airway inflammation [44], which will need further exploration. Given the critical role of CD4^+^ and CD8^+^ T cells in protection against *Mtb* [1], it was rather surprising to discover that BCG-vaccinated IRG1^-/-^ mice were as resistant as WT mice to a challenge with *Mtb* H37Rv and further support the findings of a recent study showing the importance of the itaconate pathway in linking innate immune tolerance and trained immunity [45].

On the subcellular level, we could show that, similarly to what is strongly established upon LPS/IFNγ stimulation [9, 10, 21], *Irg1* expression is highly induced during *Mtb* infection, which is a transient process due to the demonstrated negative feedback interaction between IRG1 expression and activity of the E3 ligase A20 [46, 47]. Furthermore, mycobacterial infections activate IRF1 nuclear translocation and the expression of IRG1 Importantly, in our study we could not only show that MCVs are in close vicinity of mitochondria, but also that IRG1 is recruited to MCVs, as it has been demonstrated previously for vacuoles containing *Legionella pneumophila* [23] and *Salmonella enterica* [28]. The role of direct IRG1 recruitment to pathogenic vacuoles still remains elusive, but one important molecular player in this interaction appears to be Rab32. This GTPase has been identified in a cell-intrinsic host defense mechanism able to restrict the replication of intravacuolar pathogens [48, 49], which has been shown to require IRG1 interaction to facilitate the delivery of itaconate to *Salmonella*-containing vacuoles [28]. Whether a similar mechanism is engaged during IRG1 recruitment to MCVs awaits further investigation. However, a polymorphism in Rab32 has been identified and was associated with increased susceptibility to *Mycobacterium leprae* infection [50], which might also affect other mycobacterial infections. IRG1 expression has been shown to be controlled by IRF1 [51], which recently has been demonstrated during *Mycobacterium avium* infection of human macrophages to induce nuclear translocation of IRF1 to induce IRG1 expression [52]. Though the authors did not detect a direct recruitment of IRG1 to vacuoles containing *M. avium*, they were able to see mitochondria in close vicinity of those MCVs and proposed a directed delivery of itaconate to MCVs as a plausible scenario.

Importantly, our *in vitro* experiments in *Mtb*-infected BMDMs and BMDCs could show that the recruitment of IRG1 to MCVs is correlated to decreased mycobacterial replication and lower numbers of LDs, while in IRG1^-/-^ phagocytes increased *Mtb* loads and uncontrolled generation of LDs were evident. A previous study showed that LD formation is resulting from immune activation of macrophages as part of their host defense mechanism against *Mtb* infection and is not only induced by the pathogen itself [53]. The authors showed that this HIF1α-dependent signaling pathway was required for the majority of LD formation in the lungs of *Mtb*-infected mice. On the other hand, *Mtb* is able to counteract fatty acid oxidation via HIF-1α activation to stimulate foamy macrophage generation [54], a nutrient-rich reservoir shown to be important for *Mtb* persistence. Previous work also showed that *Mtb*-containing vacuoles are able to migrate towards host LDs, and that oxygenated mycolic acids of *Mtb* are contributing to the foamy phenotype of those macrophages [55]. Our findings suggest that IRG1 induction by the host upon *Mtb* infection restricts LD formation to minimize mycobacterial growth, while increased LD generation in IRG1^-/-^ phagocytes favors *Mtb* replication indicating a dependence of this pathogen on host lipids stored in LDs (**Figure S5**).

Changes in available host lipids also have direct consequences on the modulation of adaptive immune responses. For example, LD generation was correlated to efficacy and regulation of cross-presentation pathways [56], which include exogenous antigens derived from *Mtb* infection. Our observations that IRG1 deficiency induced a severe reduction in T and B cell compartments in *Mtb*-infected lungs and spleens, further emphasizes the role of players of adaptive immunity in the progression of *Mtb* immunopathologies. Overall, we could show in our study that immunometabolic host responses during *Mtb* infection are essential to control infection outcome and immunopathology. IRG1 expression and itaconate production are one of the key nodes that determine efficient host immunity in TB suggesting that their modulation might be used in the future development of HDTs to improve immunometabolic host responses to *Mtb* infection.

## Material and Methods

### Mice

C57BL/6NJ wild type mice and C57BL/6NJ-*Acod1^em1(IMPC)J^*/J (IRG1^-/-^) mice deficient in *Irg1* expression were purchased from The Jackson Laboratory (Bar Harbor, ME, USA). All mice were maintained and breeding was performed in the animal facility of the Pasteur Institute of Lille, France (agreement B59-350009). All experimental procedures received ethical approval by the French Committee on Animal Experimentation and the Ministry of Education and Research (APAFIS#10232-2017061411305485 v6, approved on 14/09/2018). All experiments were performed in accordance with relevant guidelines and regulations.

### Murine bone marrow-derived macrophages (BMDM) and dendritic cells (BMDC)

Murine bone-marrow progenitors were obtained by sampling tibias and femur bones from 7 to 12 week-old C57BL/6NJ wild type and IRG1^-/-^ mice. BMDM were obtained by seeding 10^7^ bone marrow cells in 75 cm^2^ flasks in RPMI 1640 Glutamax medium (ThermoFisher Scientific) supplemented with 10% heat-inactivated Fetal Bovine Serum (FBS, ThermoFisher Scientific) and 10% L929 cell supernatant containing Macrophage Colony-Stimulating Factor (M-CSF). After 7 days incubation at 37°C in an atmosphere containing 5% CO_2_, the BMDM monolayer was rinsed with D-PBS and cells harvested with Versene (ThermoFisher Scientific). BMDC were differentiated as previously described [40]. Briefly, 2×10^7^ murine bone marrow progenitors were seeded in 100 ml RPMI-FBS supplemented with 10% J558-conditioned medium containing Granulocyte-Macrophage Colony-Stimulating Factor (GM-CSF) in 500 cm^2^ square petri dishes (Nunc). Cells were incubated at 37°C in 5% CO_2_. Fresh medium was added every 3-4 days. On day 10, the supernatant was discarded and adherent cells were harvested using DPBS containing 2 mM EDTA (Sigma-Aldrich). BMDM and BMDC were resuspended into corresponding culture medium to be used for subsequent assays.

### Bacteria

For *in vivo* studies, *Mycobacterium bovis* BCG (strain 1173P2), *Mtb* H37Ra and *Mtb* H37Rv WT strain were grown in Middlebrook 7H9 medium, as described previously [57] [40]. Recombinant strains of *Mtb* H37Rv expressing an enhanced green fluorescent protein (GFP) or a red fluorescent protein DsRed [40] were cultured in Middlebrook 7H9 medium (Difco) supplemented with 10% oleic acid-albumin-dextrose-catalase (OADC, Difco), 0.2% glycerol (Euromedex), 0.05% Tween 80 (Sigma-Aldrich) and 50 µg/ml hygromycin (ThermoFisher Scientific) or 25 µg/ml kanamycin (Sigma-Aldrich) for H37Rv-GFP or H37Rv-DsRed, respectively. Cultures were maintained for 14 days until the exponential phase was reached. Before cell infection, bacilli were washed with Dulbecco’s Phosphate Buffered Saline (DPBS, free from MgCl_2_ and CaCl_2_, ThermoFisher Scientific), resuspended in 10 mL RPMI-FBS and centrifuged at 1000 RPM for 2 min at room temperature to remove bacterial aggregates. Bacterial titer of the suspension was determined by measuring the optical density (OD_600 nm_) and GFP or DsRed fluorescence on a Victor Multilabel Counter (Perkin Elmer). The bacterial suspension was diluted at the required titer in RPMI-FBS.

### Chemicals, Dyes and Antibodies

100 ng/ml of ultrapure LPS from *E. coli* 0111:B4 (Invivogen, France) and 20 ng/ml of recombinant mouse IFNγ (ImmunoTools GmbH, Germany) were used to activate BMDMs. Polyclonal anti-*Mycobacterium tuberculosis* LAM (antiserum, Rabbit), NR-13821, was obtained through BEI Resources, NIAID, NIH, USA. Antibodies against IRG1 (Abcam ab122624) and LAMP-1 (BD Biosciences cat. 553792) were used as primary antibodies. Alexa Fluor 488 and 647 secondary antibodies, Hoechst 33342 and LipidTox Deep Red were all obtained from ThermoFisher Scientific (USA). DAPI was purchased from Sigma-Aldrich (USA).

### Infection of mice and determination of bacterial burden

Eight to twelve-week-old C57BL/6NJ WT and IRG1^-/-^ mice (n=16 per group) were inoculated with *Mtb* H37Rv (or PBS for control mice) *via* the intranasal (i.n) route (10^5^ CFU/20 µl) as described [58]. After infection, individual body weight and survival of mice were monitored. At indicated time post infection, mice were euthanized and targeted organs (lungs, spleen, liver and draining bronchial lymph node) were harvested for bacterial burden evaluation by colony forming units (CFU) enumeration. Organs were homogenized for 20 min in a tube containing 2.5 mm diameter glass beads and 1 ml of PBS using the MM 400 mixer mill (Retsch GmbH, Haan, Germany). Ten-fold serial dilutions (from 10^-2^ to 10^-9^) of each sample were plated onto 7H11 medium agar plate (Difco) supplemented with 10% oleic acid-albumin-dextrose-catalase (OADC, Difco). After a three-week growth period at 37°C, CFUs were determined at the appropriate dilution allowing optimal colonies enumeration.

### Lung histopathology

As described previously [59], mice were euthanized, lungs were perfused and fixed overnight at 4°C with 10 % neutral buffered Formalin solution (Sigma-Aldrich) before being embedded in paraffin. Tissues were sliced with a microtome and 5 µm sections were labelled by hematoxylin phloxine saffron (HPS) stain or Ziehl-Neelsen (acid-fast) stain and were examined by light microscopy for anatomopathology, as described previously [58].

### Histology by immunofluorescence

As described previously [59], lungs were perfused and fixed overnight at 4°C with 10 % neutral buffered Formalin solution (Sigma-Aldrich), washed in PBS, and incubated overnight at RT in a 20 % PBS-sucrose solution. Tissues were then embedded in the Tissue-Tek OCT compound (Sakura), frozen in liquid nitrogen, and cryostat sections (10 μm) were prepared. For staining, tissue sections were rehydrated in PBS and incubated in a PBS solution containing 1% blocking reagent (PBS-BR, Boeringer) for 20 min before incubation overnight at 4°C in PBS-BR containing any of the following mAbs or reagents: DAPI nucleic acid (Molecular Probes), phalloidin Alexa fluor 488 (Molecular Probes), Allophycocyanin-coupled BM8 (anti-F4/80, Abcam), Fluorescein-coupled HL3 (anti-CD11c, BD Biosciences), Fluorescien-coupled 145-2C11 (anti-CD3, BD Biosciences), Phycoerythrin-coupled E50-2440 (anti-Siglec-F, BD Biosciences), Alexa fluor 647-coupled 1A8 (anti-Ly6G, Biolegend), Allopphycocyanin-coupled RA3-6B2 (anti-CD45/B220, BD Biosciences). Polyclonal anti-*Mtb* LAM (antiserum, Rabbit), NR-13821, was obtained through BEI Resources, NIAID, NIH, USA. 2 hours at RT incubation with Alexa fluor 568-coupled Donkey anti-Rabbit IgG (ThermoFisher Scientific) was added to reveal anti-*Mtb* LAM staining. Slides were mounted in Fluoro-Gel medium (Electron Microscopy Sciences). Labeled tissue sections were visualized with an Axiovert M200 inverted microscope (Zeiss, Iena, Germany) equipped with a high-resolution mono-chrome camera (AxioCam HR, Zeiss). At least three slides were analyzed per organ from three different animals.

### Flow cytometry

Harvested lungs and spleens were cut into small pieces and incubated for 1 hour at 37 °C with a mix of DNAse I (100 μg/ml, Sigma-Aldrich) and collagenase D (400 U/ml, Roche). Lung cells were washed and filtered before being incubated with saturating doses of purified 2.4G2 (anti-mouse Fc receptor, ATCC) in 200 μl PBS, 0.2% BSA, 0.02% NaN_3_ (FACS buffer) for 20 min at 4 °C. Various fluorescent monoclonal antibody combinations in FACS buffer were used to stain cells. Acquisitions were done on a FACS Canto II flow cytometer (Becton Dickinson) with the following mAbs: Fluorescein (FITC)-coupled 145-2C11 (anti-CD3, BD Biosciences), FITC-coupled HL3 (anti-CD11c, ThermoFisher Scientific), Phycoerythrine (PE)-coupled M1/70 (anti-CD11b, BD Biosciences), PE-coupled 53-6.7 (anti-CD8a, BD Biosciences), Allophycocyanin (APC)-coupled BM8 (anti-F4/80, BD Biosciences), APC-coupled RM4-5 (anti-CD4, Biolegend), Brillant violet 421 (BV421)-coupled E50-2440 (anti-SiglecF, BD Biosciences), BV421-coupled M5 (anti-MHCII, BD Biosciences), PE/cyanine(Cy)7-coupled RA3-6B2 (anti-CD45/B220, BD Biosciences) and PE/Cy7-coupled 1A8 (anti-Ly6G, Biolegend). Fixable viability dye Aqua (ThermoFisher Scientific) was used to gate viable cells.

| Cell type | Phenotype |
| --- | --- |
| Neutrophils | CD11b<sup>+</sup> Ly6G<sup>+</sup> |
| Dendritic cells | CD11b<sup>+</sup> CD11c<sup>+</sup> F4/80<sup>-</sup> |
| Alveolar macrophages | F4/80<sup>+</sup> SiglecF<sup>+</sup> CD11c<sup>+</sup> |
| Interstitial macrophages | F4/80<sup>+</sup> SiglecF<sup>-</sup> CD11c<sup>int</sup> |
| Eosinophils | F4/80<sup>+</sup> SiglecF<sup>+</sup> CD11c<sup>-</sup> |
| CD4 T cells | CD3<sup>+</sup> CD4<sup>+</sup> |
| CD8 T cells | CD3<sup>+</sup> CD8<sup>+</sup> |
| B cells | B220<sup>+</sup> MHCII<sup>+</sup> |

### Quantitative RT-PCR

BMDMs were grown in 6-well plates and RNA was isolated using the RNeasy Mini kit (Qiagen) following the manufacturer’s instructions. RNA concentration was determined using the GE SimpliNano device (GE Healthcare). Remaining DNA in samples was digested using the amplification grade DNase I kit (Sigma-Aldrich) for 6 min at RT. The reaction was stopped by heat inactivation for 10 min at 70°C. cDNA synthesis was achieved by reverse transcription using the SuperScript VILO kit (ThermoFisher Scientific) following the manufacturer’s instructions. qPCR was performed using the LightCycler 480 SYBR Green I reagent (Roche) with 20 ng cDNA per sample and the indicated primer pairs (**Supplemental Table S1**). qPCR reactions were measured by the QuantStudio 12K Flex system (Applied Biosystems) using the following cycles: 2 min 50°C, 10 min 95°C followed by 40 cycles of 15 s at 95°C, 30 s at 60°C and 30 s at 72°C.

### Infection for intracellular mycobacterial replication and lipid droplet (LD) formation assays

2×10^4^ BMDM or 4×10^4^ BMDC were seeded per well in 384-well plates (µClear, Greiner Bio-One). After 12 hours incubation at 37°C with 5% CO_2_, LPS (100 ng/mL) and IFN-gamma (50 ng/mL) were added as positive control. After overnight incubation, cells were infected for 2 h with H37Rv-GFP at a MOI of 1. Cells were washed with RPMI-FBS and treated with amikacin (50 µg/mL) for 1 h in order to remove extracellular *Mtb*. Then, cells were washed twice with RPMI-FBS and incubated at 37°C with 5% CO_2_.

For intracellular mycobacterial replication assay, 10% formalin solution (Sigma-Aldrich) containing 10 µg/mL Hoechst 33342 (ThermoFisher Scientific) was replaced to each well at 2 h and 96 h post-infection. Plates were incubated at RT for 30 min, allowing staining and cell fixation. Cells were stored in DPBS supplemented with 1% FBS, until image acquisition.

For LD formation assay, cells were washed and fixed at 96 h post-infection, as previously described [40]. Cells were washed twice with DBPS and intracellular LDs were stained with 25 µL per well of 2000-fold diluted HCS LipidTOX deep Red neutral lipid probe (ThermoFisher Scientific) in PBS for 30 min at room temperature.

### Image Acquisition

Image acquisitions were performed on an automated confocal microscope (Opera QEHS, PerkinElmer) using a 20x and 60x water objectives for intracellular mycobacterial replication and LD formation assays, respectively. The microscope was equipped with 405 nm, 488 nm, 561 nm and 640 nm excitation lasers. The emitted fluorescence was captured using three cameras associated with a set of filters covering a detection wavelength ranging from 450 nm to 690 nm. Hoechst 33342-stained nuclei were detected using the 405 nm laser with a 450/50-nm emission filter. Green or red signals, corresponding to H37Rv-GFP and H37Rv-DsRed, were recorded using 488 nm or 561 nm lasers with 540/75- or 600/40-nm emission filters, respectively. LipidTOX signal was detected using 630-nm excitation and 690-nm emission wavelengths.

### Image-based analysis

For mycobacteria replication and LD generation quantification, images from the automated confocal microscope were analyzed using multi-parameter scripts developed using Columbus system (version 2.3.1; PerkinElmer) as described previously [40] (**Supplemental Tables 2 and 3**).

#### Cell detection and M. tuberculosis intracellular replication

Segmentation algorithms were applied to input images to detect nuclei labeled by Hoechst 33342 (cyan) and the GFP signal of *Mtb* H37Rv (green) to determine infection and replication rates. Briefly, the host cell segmentation was performed using two different Hoechst signal intensities—a strong intensity corresponding to the nucleus and a weak intensity in cytoplasm—with the algorithm “Find Nuclei” and “Find Cytoplasm”, as described previously [60]. GFP or DsRed signal intensities in a cell were used for the intracellular bacterial segmentation with the algorithm “Find Spots”. The identified intracellular bacteria were quantified as intracellular *Mtb* area with number of pixels. Subsequently, population of infected cells was determined, and the increase of intracellular *Mtb* area, corresponding to intracellular mycobacterial replication, was calculated.

#### Cell detection and quantification of LD formation

LDs labeled by LipidTox DeepRed (red), nuclei labeled by Hoechst 33342 (blue) and the GFP signal of *Mtb* H37Rv (green) were detected using segmentation algorithms applied to input image. Briefly, the host cell segmentation using Hoechst signal and LipidTox intensities was performed to detect nuclei and cell borders, respectively. *Mtb* and LDs were determined by applying masks, which were adapted to GFP and LipidTox dye signal intensities, respectively. Further filtering and refinement using the algorithm “Find Micronuclei” and based on size-to-signal intensity and area of LD candidates allowed specific selection of LDs, which were separated from out-of-focus and background signal intensities. The identified intracellular bacteria were quantified as intracellular *Mtb* area with number of pixels. Subsequently, population of infected (*Mtb*) and non-infected (NI) cells were determined, and the percentage of LD-positive cells and the average of LD number per cell were calculated.

### Immunofluorescence

BMDMs were grown overnight on poly-L-lysine-coated glass coverslips the day before the infection. Cells were infected with *Mtb* H37Rv-DsRed at an MOI = 1 for 24 h. Subsequently, cells were washed three times with PBS and fixed in PBS + 4% paraformaldehyde + 4% sucrose, pH 7.4, for 20 min at RT followed by quenching in PBS + 50 mM NH_4_Cl for 10 min. Coverslips were labeled for IRG1 (Abcam, UK), DNA using DAPI (Sigma-Aldrich, USA) and F-actin using Alexa Fluor 680 phalloidin (ThermoFisher Scientific). After extensive washing, the coverslips were post-fixed in 4% paraformaldehyde for 16 h and then mounted with ProLong Gold antifade reagent (ThermoFisher Scientific). Images were acquired using a confocal microscope (Zeiss LSM880) equipped with a 63x objective (NA 1.4) and Zen imaging software (Zeiss, Germany).

### Processing for conventional electronic microscopy

Infected BMDMs were fixed 4 days post-infection at room temperature for 1 h with 2.5% glutaraldehyde in Na-cacodylate buffer 0.1 M (pH 7.2) containing 0.1 M sucrose, 5 mM CaCl_2_, and 5 mM MgCl_2_, washed with complete cacodylate buffer and post-fixed for 1 h at room temperature with 1% osmium tetroxide in the same buffer devoid of sucrose. Cells were washed with the same buffer, scraped off the dishes and concentrated in 2% agar in Na-cacodylate buffer. Subsequently, samples were washed twice in Na-cacodylate buffer and dehydrated in a graded series of aceton solutions and gradually incorporated in Spurr resin. Ultrathin sections (80 nm) were cut with an ultracryomicrotome (EM UC7, Leica) and were stained with 1% uranyl acetate in ultrapure water and then with Reynolds reagent. Samples were analyzed with a Tecnai G2 20 TWIN (200 kV) transmission electron microscope (FEI) and digital images were acquired with a digital camera (Eagle, FEI) for further quantification. For image analysis, precautions were taken to avoid analysis of serial cuts, and 100 WT BMDMs and 64 IRG1^-/-^ BMDMs were examined randomly by TEM to determine the number of bacteria and number of lipid granules per cell.

### Western Blotting

Latex bead-containing phagosomes (LBPs) were isolated as described previously [22]. LBP pellets were lysed in 2% (v/v) Triton X-100, 50 mM Tris, pH 8.0, 10 mM dithiothreitol, 2x protease inhibitor mixture (Roche, France) for 30 min on ice. Phagosomal lysates and total cell lysates (TCL) were mixed with 5x Laemmli sample buffer and boiled for 5 min at 95 °C. Samples were loaded on 4-15% Criterion TGX protein gels (Bio-Rad) and run in 25 mM Tris, 192 mM glycine, 0.1% (m/vol) SDS, pH 8.3, at 100 V. Proteins were transferred onto 0.2 µm PVDF membranes by a Trans-Blot Turbo device (Bio-Rad) at 2.5 A for 7 min. Equal loading of samples was controlled by Ponceau S staining. After transfer, membranes were blocked in 5% dry milk and incubated with primary antibodies and peroxidase-conjugated secondary antibodies. Bound antibodies were revealed using Chemiluminescence Blotting Substrate from Roche according to the manufacturers’ suggestions.

### Microscale oxygraphy

Real-time extracellular acidification rate (ECAR) and oxygen consumption rate (OCR) were measured using the Seahorse XFe24 flux analyzer (Seahorse Bioscience, Agilent) as described previously [61]. Briefly, BMDMs were seeded at a density of 1.6×10^6^ cells/ml the day before the measurement and incubated at 37°C and 5% CO_2_. BMDMs were treated or not with 100 ng/ml of ultrapure LPS and 20 ng/ml of recombinant mouse IFNγ for 18 h. On the day of the measurement, culture medium was removed and cells were washed once with assay medium consisting of DMEM (Sigma-Aldrich), pH buffered at 7.35, supplemented with 10 mM glucose and 2 mM glutamine and were incubated at 37 °C for 30 min. After baseline measurements, ECAR was analyzed after injections of D-glucose (10 mM), oligomycin A (1 μM), an ATP synthase inhibitor, and 2-deoxyglucose (10 mM), a competitive inhibitor of glucose. Similarly, after baseline measurements, OCR was analyzed after subsequent injection of oligomycin A (1 μM), two injections of carbonyl cyanide-4 (trifluoromethoxy) phenylhydrazone (FCCP, 0.27 μM and then 0.34 μM), a protonophore, and injection of a rotenone/antimycin A mix (1 μM), inhibitors of complex III and I. Mitochondrial spare reserve capacity (SRC) was calculated by subtracting basal respiration from maximal respiratory capacity as described previously [36].

### Statistics

All analyses and histograms were performed using GraphPad Prism 7 software. Significance of obtained results was tested using Mantel-Cox test (survival of *Mtb*-infected mice) and by Mann-Whitney test (Plating of CFU of different organs, immune cell profiling). Differences in the mean between two groups were analyzed using Student’s t-test (*in vitro* replication and LD experiments). Indicated symbols of *, ** and *** denote p < 0.05, p < 0.01 and p < 0.001, respectively.

## Author contributions

A.M., P.B., and E.H. conceived the study and wrote the manuscript. A.M., I.B., N.D., I.P., J-P.S-A., A-M.P., O-R.S., S.J., C.R., A.P., S.M., W.L., J.K., and S.C. performed and analyzed experiments. J-P.S-A., C.R., J.K., E.M., R.B., and S.C. provided reagents. S.M. and L.M. provided expertise and feedback on the study. P.B. and E.H. secured funding.

## Acknowledgments

We gratefully acknowledge Alexandre Vandeputte, Gerald Larrouy-Maumus, Ruxandra Gref, Tom Bourguignon, Alain Baulard, Cyril Gaudin, Muriel Pichavant, Isabelle Ricard, and Martin Cohen-Gonsaud for technical assistance and helpful discussions. We thank Nicolas Vandenabeele and Robin Prath for expert BSL-3 animal facility assistance. We acknowledge the PICT-IBiSA, Elisabeth Werkmeister, Hélène Bauderlique and Frank Lafont from BiCEL for providing access to microscopy and flow cytometry equipment. Financial support for this work was provided by the European Community (CycloNHit n° 608407, ERC-STG INTRACELLTB n° 260901), the Agence Nationale de la Recherche (ANR-10-EQPX-04-01, ANR-14-CE08-0017, ANR-16-CE35-0009, ANR-18-JAM2-0002, ANR-20-CE44-0019), the EMBO Young Investigator Program, the Feder (12001407 (D-AL) Equipex Imaginex BioMed), the I-SITE ULNE Foundation (ERC Generator Grant) and the Fondation pour la Recherche Medicale (SPF20170938709). The funders had no role in study design, data collection and analysis, decision to publish, or preparation of the manuscript.

## Supplemental figures

**Figure S1.**
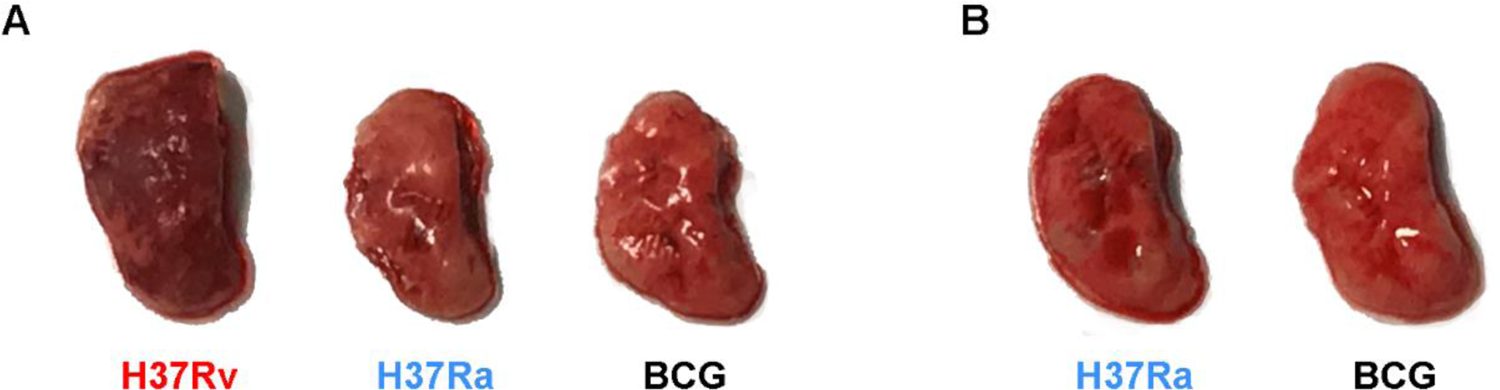
Mouse lungs infected by different mycobacterial strains at 84 dpi. WT mice **(A)** and of IRG1^-/-^ mice **(B)** were infected intranasally as indicated in Fig. 1A with the indicated mycobacterial strains and their lungs analyzed for pathological differences at 84 dpi.

**Figure S2.**
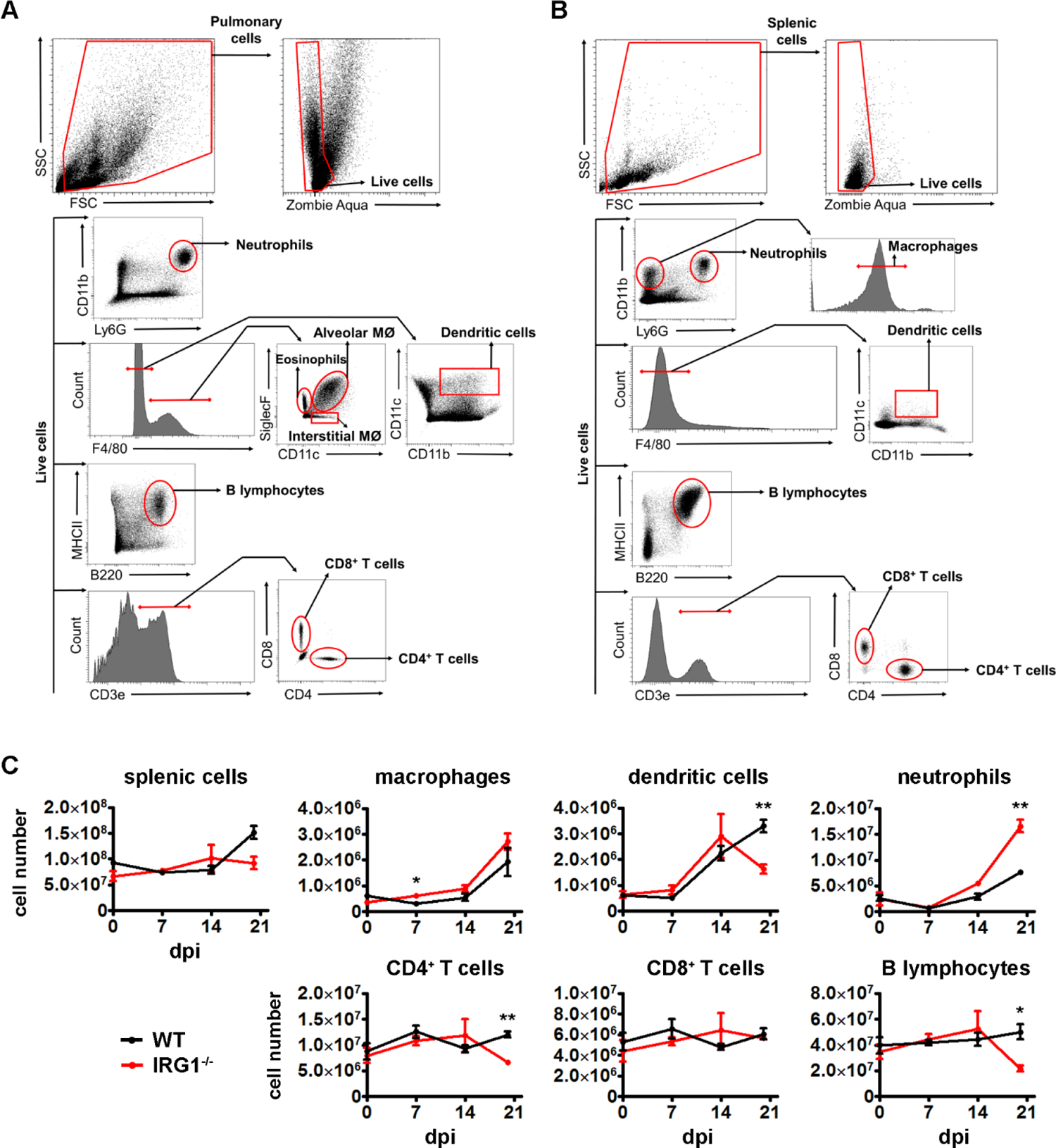
Flow cytometry analysis of pulmonary and splenic subpopulations of infected mice. Flow cytometry gating strategies for pulmonary and splenic cell populations of interest. Cells were first identified using forward scatter (FSC) and side scatter (SSC) gates to exclude residual red blood cells and cellular fragments. Among these, alive cells were selected using a fixable viability dye (Zombie Aqua). Whole organ cell tissues were stained to identify different immune cell types. **(A)** In lungs, neutrophils were selected after gating CD11b+ and Ly6G+ cells (CD11b+ Ly6G+), while classical dendritic cells were identified using positive selection of CD11b and CD11c markers on cells negative for F4/80 (F4/80-CD11b+ CD11c+). On positive F4/80 cells, SiglecF and CD11c markers were used to discriminate eosinophils (F4/80+ SiglecF+ CD11c-), alveolar macrophages (MØ) (F4/80+ SiglecF+ CD11c+) and interstitials MØ (F4/80+ SiglecF-CD11c^int^). B cells were identified using MHCII and CD45/B220 markers (MHCII+ B220+). CD3, CD4 and CD8 markers were used to select CD4+ T cells (CD3+ CD8-CD4+) and CD8+ T cells (CD3+ CD8+ CD4-), respectively. **(B)** In spleens, similar gating strategies were used to select neutrophils, classical dendritic cells, B lymphocytes, CD4 T cells and CD8 T cells. Macrophages were discriminated by the selection of F4/80+ cells among the CD11b+ Ly6G-cell population (CD11b+ Ly6G-F4/80+). **(C)** Profiling of splenic immune cells of WT mice and IRG1^-/-^ mice during the course of *Mtb* H37Rv infection. Histograms depict changes in cell numbers of the indicated immune cell populations, as determined by flow cytometry using specific cell surface markers. All cell numbers were normalized to the total cell number analyzed in each sample and were extrapolated to the whole organ. Shown are mean ± SEM of cells obtained from three individual mice per group. Statistical differences were determined by non-parametric Mann-Whitney test (* P value < 0.05, ** P value < 0.01).

**Figure S3.**
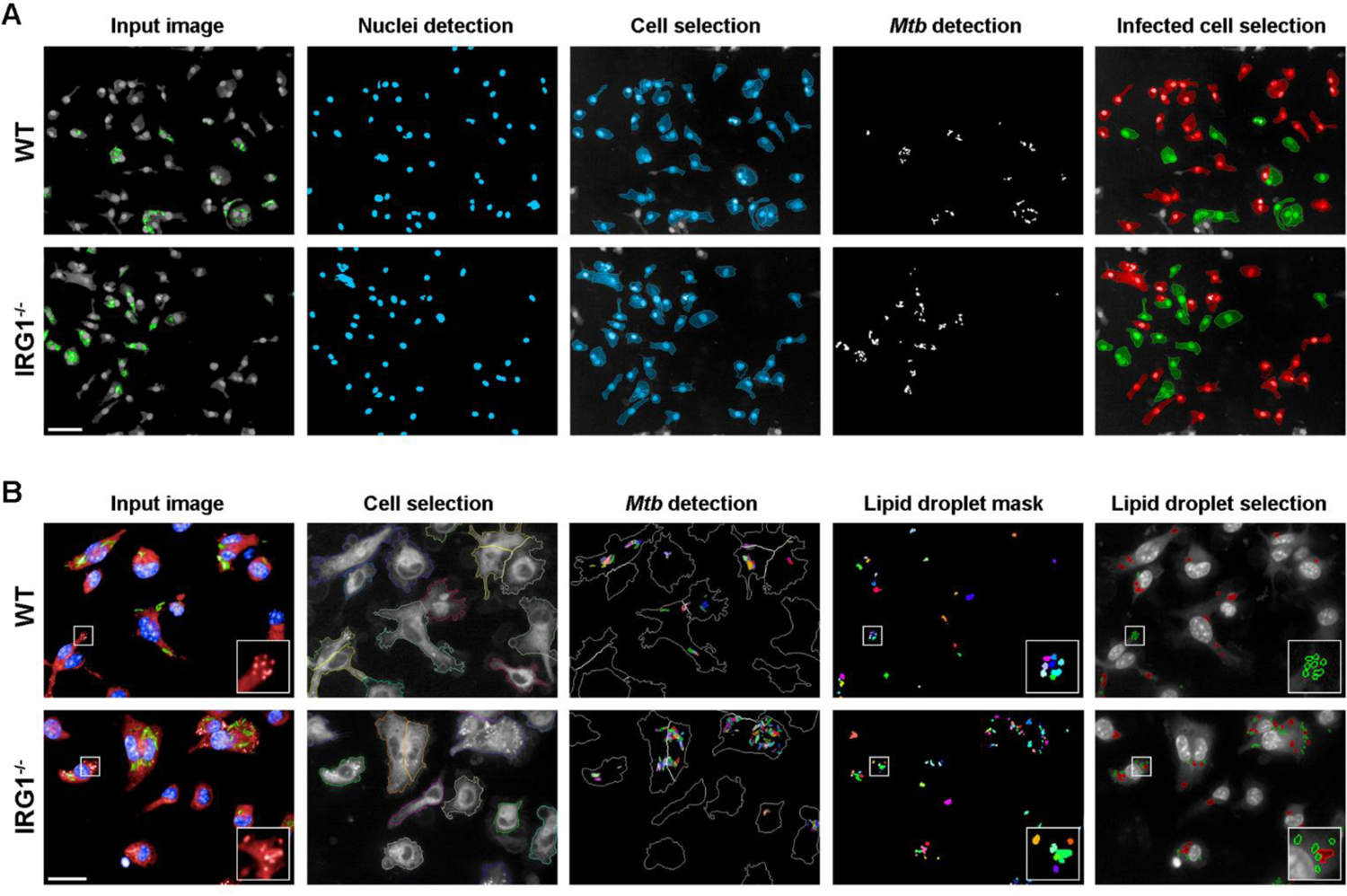
Workflows of the applied automated, multiparametric image analysis pipeline using Columbus software. **(A)** Workflow to assess the intracellular replication of *Mtb* in BMDMs and BMDCs. Segmentation algorithms were applied to input images of WT cells (upper panel) and IRG1^-/-^ cells (lower panel) to detect nuclei and cell borders (labeled by Hoechst 33342) and to identify *Mtb* H37Rv (expressing GFP) to determine infection and replication rates. In the right panel, infected cells are depicted in green, while non-infected cells are shown in red. Cells only partially shown in the microscopy field (depicted in gray) were excluded from the analysis. Bar: 50 µm. **(B)** Workflow to quantify the generation of lipid droplets (LDs). BMDMs and BMDCs were labeled with LipidTox DeepRed to visualize LDs (red), nuclei were labeled by Hoechst 33342 (blue) and the GFP signal of *Mtb* H37Rv (green). Segmentation algorithms were used to detect cell borders and *Mtb* followed by an LD mask adapted to LipidTox dye signal intensities. Further filtering and refinement based on size-to-signal intensity allowed specific selection of LDs (green circles in the right images), which were separated from out-of-focus and background signal intensities (red circles in the right image panel). Insets show magnifications of LDs of example cells. Bar: 20 µm. Further details of the applied scripts are indicated in Supplemental tables 2 and 3.

**Figure S4.**
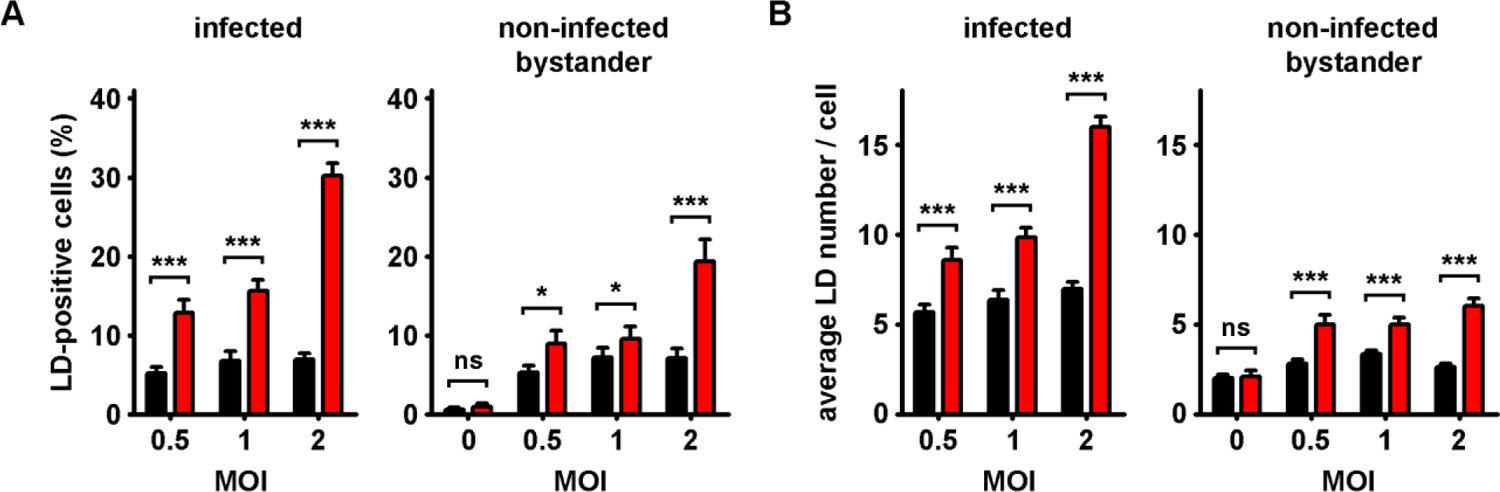
IRG1 deficiency leads to increased amounts of lipid droplets (LDs) in infected as well as non-infected bystander phagocytes. **(A)** Histograms depicting the percentage of LD-positive, *Mtb*-infected BMDCs (left panel) and non-infected bystander BMDCs (right panel) at 96 hpi, as determined by automated confocal microscopy. **(B)** Histograms showing the average LD number per cell for infected and non-infected bystander BMDCs at 96 hpi. Shown are mean ± SEM of at least 6 analyzed wells per condition (n > 400 cells) of one representative out of three independent experiments. Statistical differences were determined by Student’s t-test (ns: non-significant, * P value < 0.05, *** P value < 0.001).

**Figure S5.**
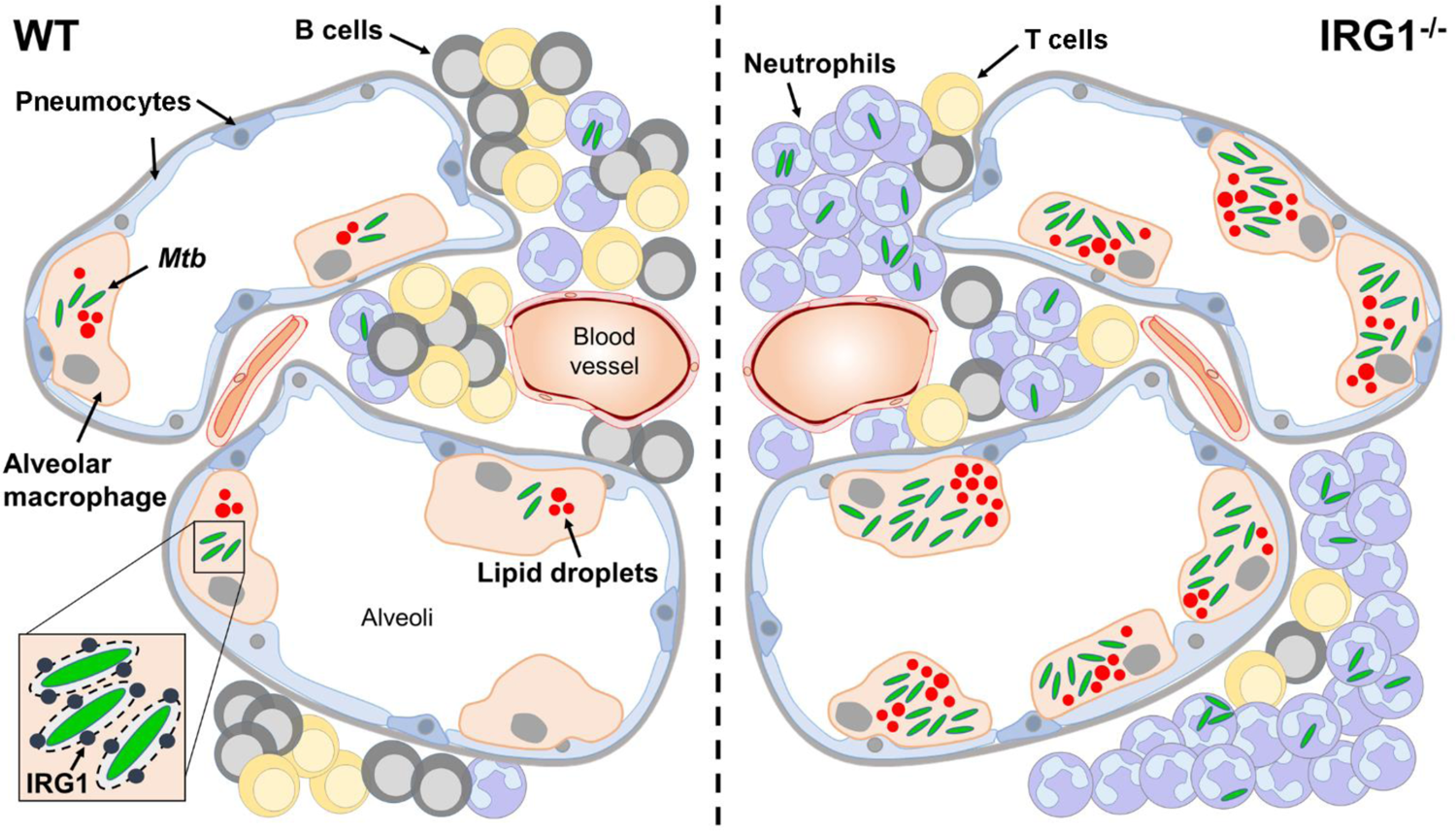
Induced expression of IRG1 during *Mtb* infection restricts formation of lipid droplets and intracellular pathogen growth. Infection of alveolar macrophages by *Mtb*, which leads to recruitment of IRG1 to *Mtb*-containing vacuoles (inset) in WT conditions (left panel), restricts generation of lipid droplets, an important nutrient reservoir of *Mtb* in host cells. Limited availability of host lipids confines intracellular replication of the pathogen and permits induction of adaptive immune responses. Absence of IRG1 during *Mtb* infection (right panel) leads to uncontrolled formation of lipid droplets and exacerbated pathogen growth in host cells. Increased inflammation in the lung and infiltration of neutrophils further allows *Mtb* dissemination and reduces numbers of recruited B and T lymphocytes, which possibly attenuate adaptive immunity.

## Supplemental tables

**Table S1.**
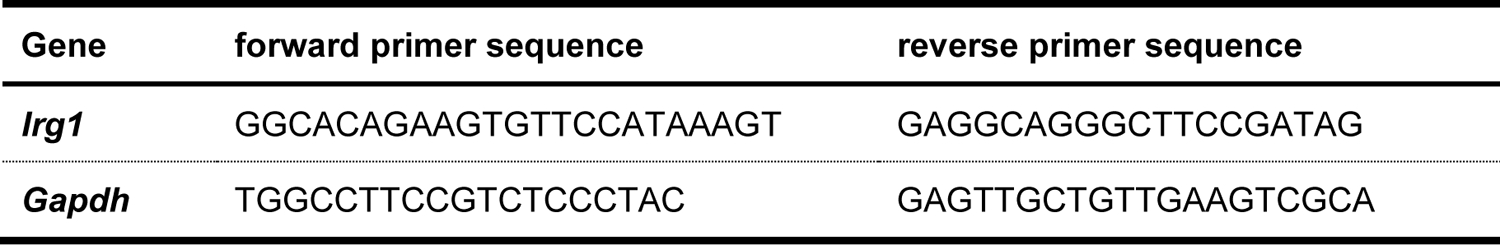
Sequences of the RT-PCR primers used in this study.

**Table S2.** Applied in-house multi-parametric script used in Columbus (PerkinElmer) to determine *Mtb* replication.

| Input Image | Stack Processing:<br>Individual Planes<br>Flat field<br>Correction: None | Method | Output |
| --- | --- | --- | --- |
| Calculate Image | | Method : By Formula<br>Formula : $\text{iif}(A > 200, A, 0)$<br>Channel A : Exp1Cam1<br>Negative Values : Set to Zero<br>Undefined Values : Set to Local Average | Output Image :<br>DAPImask |
| Find Nuclei | Channel : DAPImask<br>ROI : None | Method : B<br>Common Threshold : <u>0.8</u><br>Area : $> 30 \mu\text{m}^2$<br>Split Factor : 7<br>Individual Threshold : 0.4<br>Contrast : $> 0.1$ | Output<br>Population :<br>Nuclei |
| Find Cytoplasm | Channel :<br>Exp1Cam1<br>Nuclei : Nuclei | Method : A<br>Individual Threshold : <u>0.3</u> |  |
| Select Population | Population : Nuclei | Method : Common Filters<br>Remove Border Objects<br>Region : Cell | Output<br>Population : Cells<br>selected |
| Calculate Image | | Method : By Formula<br>Formula : $\text{iif}(A > 200, A, 0)$<br>Channel A : Exp2Cam2<br>Negative Values : Set to Zero<br>Undefined Values : Set to Local Average | Output Image :<br>Mtb Mask |
| Find Spots | Channel : Mtb Mask<br>ROI : Cells selected | Method : A<br>Relative Spot Intensity : $> 0.03$<br>Splitting Coefficient : <u>0.54</u><br>Calculate Spot Properties | Output<br>Population :<br>Intracellular Mtb |
| Calculate Morphology Properties | Population :<br>Intracellular Mtb<br>Region : Spot | Method : Standard<br>Area | Output<br>Properties :<br>Intracellular Mtb |
| Calculate Properties | Population : Cells<br>selected | Method : By Related Population<br>Related Population : Intracellular Mtb<br>Number of Intracellular Mtb<br>Intracellular Myb Area [$\text{px}^2$] | Output<br>Properties : per<br>Cell |
| Select Population | Population : Cells<br>selected | Method : Filter by Property<br>Number of Intracellular Mtb- per Cell : $> 0$ | Output<br>Population :<br>Infected cells |
| Define Results | Method: List of Outputs<br>Number of Objects<br>Population : Intracellular Mtb<br>Intracellular Mtb Area [$\text{px}^2$] : Sum<br>Population : Infected Cells<br>Intracellular Mtb Area [$\text{px}^2$]- Sum per Cell : Mean<br>Method : Formula Output<br>Formula : $a/b \times 100$<br>Population Type : Objects<br>Variable A : Infected Cells - Number of Objects<br>Variable B : Cells Selected - Number of Objects<br>Output Name : % Infected cells | | |

**Table S3.**
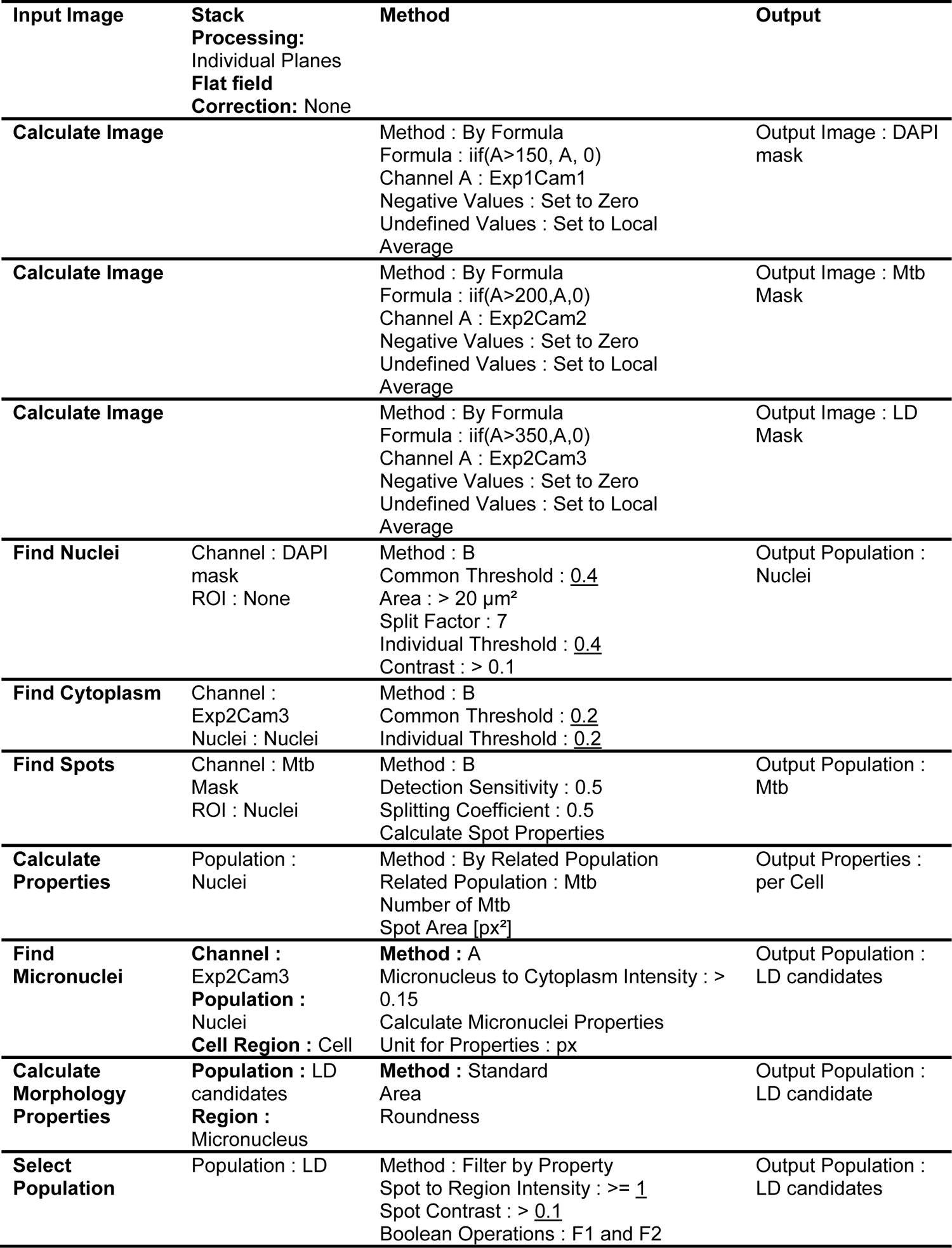
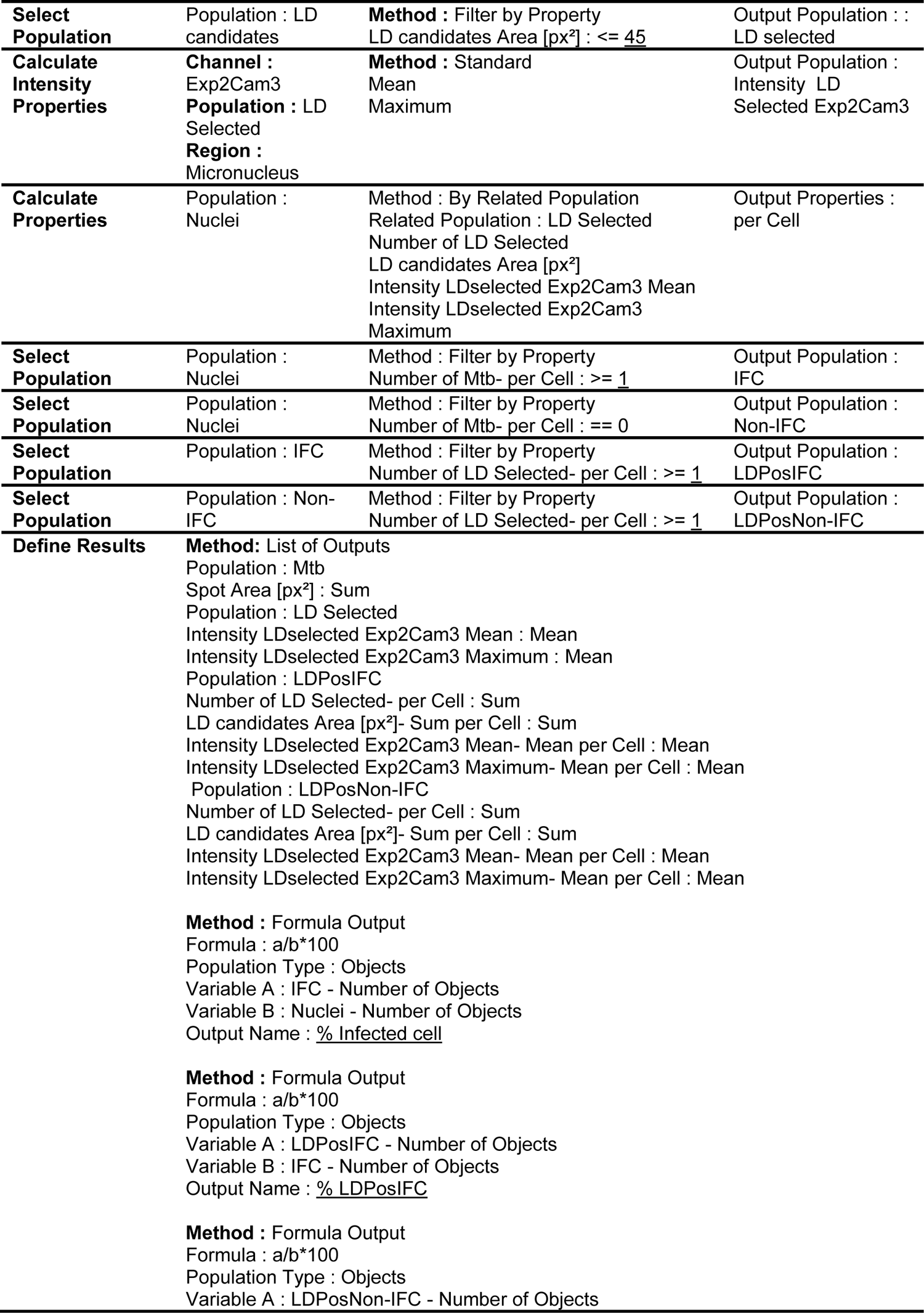
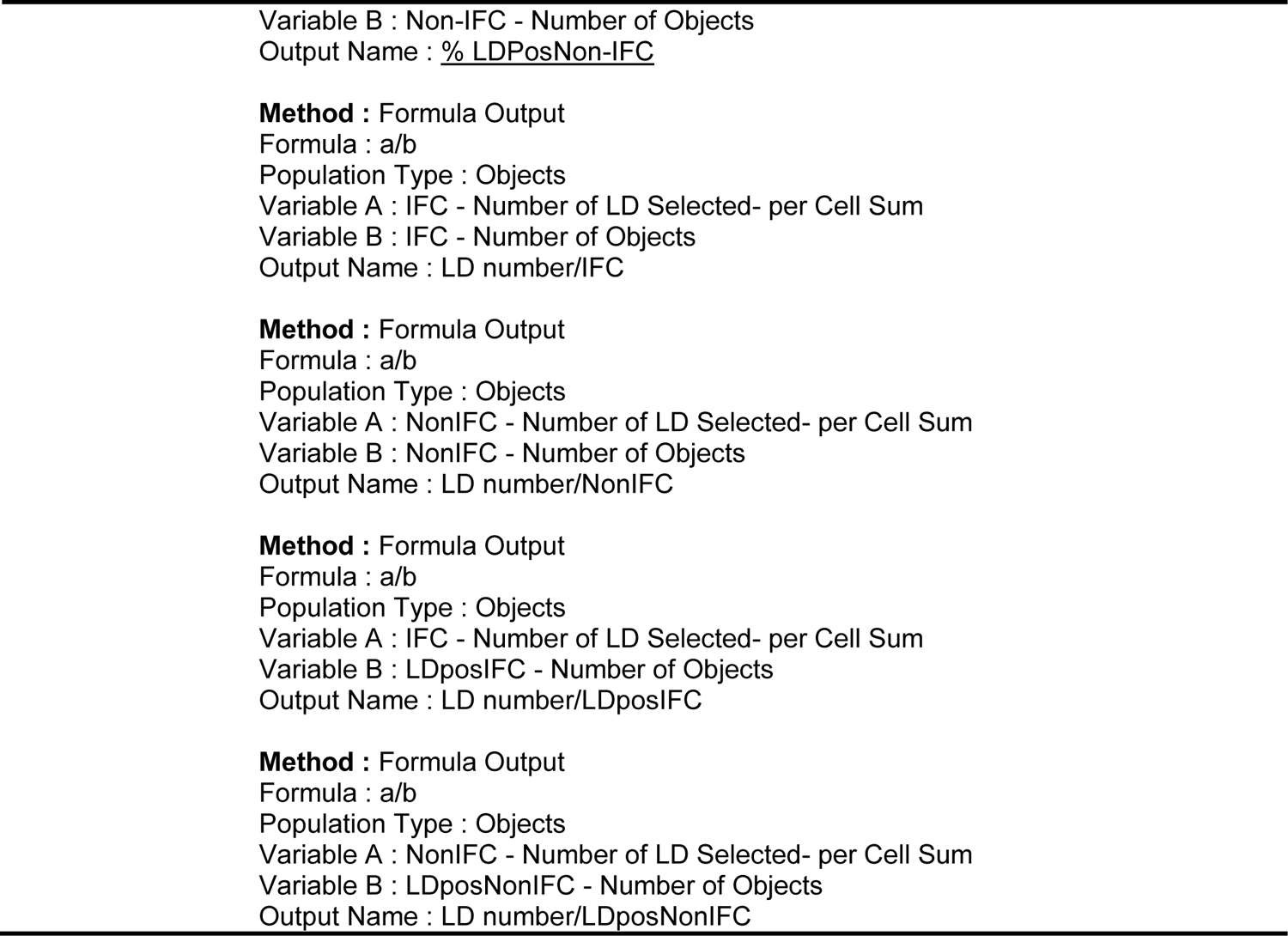
Applied in-house multi-parametric script used in Columbus (PerkinElmer) to determine lipid droplet generation.

